# Light-mediated discovery of surfaceome nanoscale organization and intercellular receptor interaction networks

**DOI:** 10.1101/2020.08.11.246652

**Authors:** Maik Müller, Fabienne Gräbnitz, Niculò Barandun, Yang Shen, Stefan U. Vetterli, Milon Mondal, James R. Prudent, Yannik Severin, Marc van Oostrum, Raphael Hofmann, Roman Sarott, Alexey I. Nesvizhskii, Erick M. Carreira, Jeffrey Bode, Berend Snijder, John A. Robinson, Martin J. Loessner, Annette Oxenius, Bernd Wollscheid

## Abstract

Delineating the molecular nanoscale organization of the surfaceome is pre-requisite for understanding cellular signaling. Technologies for mapping the spatial relationships of cell surface receptors and their extracellular signaling synapses would open up theranostic opportunities and the possibility to engineer extracellular signaling. Here, we developed an optoproteomic technology termed LUX-MS that exploits singlet oxygen generators (SOG) for the light-triggered identification of acute protein interactions on living cells. Using SOG-coupled antibodies, small molecule-drugs, biologics and intact viral particles, we show that not only ligand-receptor interactions can be decoded across organisms, but also the surfaceome receptor nanoscale organization ligands engage in with direct implications for drug action. Furthermore, investigation of functional immunosynapses revealed that intercellular signaling inbetween APCs and CD8^+^ T cells can be mapped now providing insights into T cell activation with spatiotemporal resolution. LUX-MS based decoding of surfaceome signaling architectures provides unprecedented molecular insights for the rational development of theranostic strategies.

## Introduction

Cellular function is regulated by information exchange with the outside world. The cell surface proteotype (surfaceome) is the signaling gateway to the cell that enables and limits communication^1^. Nanoscale receptor assemblies connect external stimuli with intracellular response pathways within surfaceome signaling domains^2^. These substructures play critical roles in drug action, host-pathogen interactions and intercellular communication^3^. As such, mapping ligand-targeted surfaceome landscapes with nanoscale precision should provide fundamental insights into cellular signaling function in health and disease with wide-reaching implications for the development of therapeutic strategies. Specifically, unravelling surfaceome signaling interactions in space and time would substantially advance drug development (deconvoluting drug targets and mechanism of action) and enable discoveries in broad research areas including cell biology (dissecting receptor signaling) ^4^, microbiology (identifying pathogen binding sites)^5^ and immunology (elucidating immunosynaptic communication)^6^.

Still today, the surfaceome signaling landscape remains *terra incognita* that cannot be inferred from cellular proteomes and transcriptomes. Dedicated strategies have been established to profile cellular surfaceomes^7^. For example, the extensive application of the Cell Surface Capture (CSC) technology^8,9^ established N-glycosylated surfaceomes for numerous cell types (collectively reported in the Cell Surface Protein Atlas, CSPA https://wlab.ethz.ch/cspa/)^10^ and enabled the *in silico* characterization of the entire human surfaceome^11^. While, fluorescence-based reporter assays^12–14^ and platforms of genetically engineered receptors^15–17^ established associations between receptors by high-throughput testing of binary interactions within and across surfaceomes, the Ligand-Receptor Capture (LRC) technology identified direct ligand interactions of N-glycosylated receptors^18,19^. Recently, enzyme-based proximity labeling strategies such as APEX^20^, HRP or PUP-iT^21^ enabled labeling and MS-based identification of lipid rafts^22^, growth factor signaling domains^23^, B-cell receptor neighborhoods^24^ and intact neuronal synapses^25^ on living cells and in fixed tissues^26^. A photocatalyst-based proximity labeling strategy thereby showed improved spatial precision^27^. However, in such methods, labeling probes are simultaneously allocated to all surface accessible receptors by antibodies or genetic fusion resulting in a lack of flexibility and specificity to detect ligand-targeted surfaceome signaling domains, leaving the functional hot-spots of the cellular surfaceome landscape unexplored.

Singlet oxygen generators (SOG) are known to exert proximity labeling capabilities through light-activated generation of reactive singlet oxygen species that oxidize biomolecules in nanometer vicinity^28,29^. Small molecule SOG^30^ and genetically encodable SOG are extensively used throughout life sciences for photodynamic inactivation of proteins^31–34^ and cells^35,36^, for correlative light-electron microscopy^37^ and for the detection of intracellular protein-protein interactions^38,39^ in a variety of organisms. The capability of SOG to decode cell surface interactions was recently demonstrated by identifying the binding target of a nerve-highlighting fluorescent peptide in mammalian tissues^40^. Due to the small size and light-controlled *in-situ* tagging capabilities, SOG represent ideal probes for ligand-guided labeling and detection of surfaceome signaling domains.

Here, we developed and applied a light-controlled proximity detection technology termed LUX-MS to systematically explore the surfaceome signaling landscape on living cells. The approach capitalizes on ligand-coupled small molecule SOG for the specific and light-tunable biotinylation of ligand-proximal proteins enabling their identification using a high-throughput robot-assisted proteotyping workflow. We demonstrate the spatial specificity and broad applicability of our approach to decode surfaceome signaling domains of small molecules, biomolecules and intact pathogens in mammalian and bacterial systems in a discovery-driven fashion and without the need for genetic manipulation. Lastly, we showcased the utility of LUX-MS to elucidate complex intercellular surfaceome signaling domains by mapping the molecular architecture of functional immunosynapses in unprecedented detail.

## Results

### Development of the LUX-MS technology

We first tested the capability of the small-molecule SOG thiorhodamine to photo-oxidize transferrin proteins *in vitro*. Quantitative MS analysis and open modification search revealed light-dependent conversion of the amino acids histidine, cysteine, tryptophan, and methionine into previously described photo-oxidation products^28^ (**Supplementary Fig. 1**). Among others, the most prevalently formed 2-oxo-histidine (His+14 Da) modification bears a light-induced ketone group that offers functionalization via hydrazone formation using hydrazide-containing linkers. Indeed, incubation of photo-oxidized transferrin with a biotin-hydrazide (BH) linker led to significant consumption of 2-oxo-histidine (**Supplementary Fig. 2**) indicating biotinylation of light-modified proteins. Next, we combined light-controlled protein biotinylation with liquid-handling robot-assisted processing and high-performance liquid chromatography tandem mass spectrometry (LC-MS/MS) to establish the optoproteomic LUX-MS technology (**Fig. 1**). SOG-proximal proteins are thereby biotinylated in live cells under physiological conditions by light and are captured after cell lysis using streptavidin-functionalized tips. On-bead digestion using proteolytic enzymes releases peptides N- and C-terminally of the initial labeling site that are subsequently label-free quantified by mass spectrometry. SOG-proximal proteins are eventually identified in a hypothesis-free fashion by light-dependent enrichment against a non-illuminated control.

**Fig. 1.**
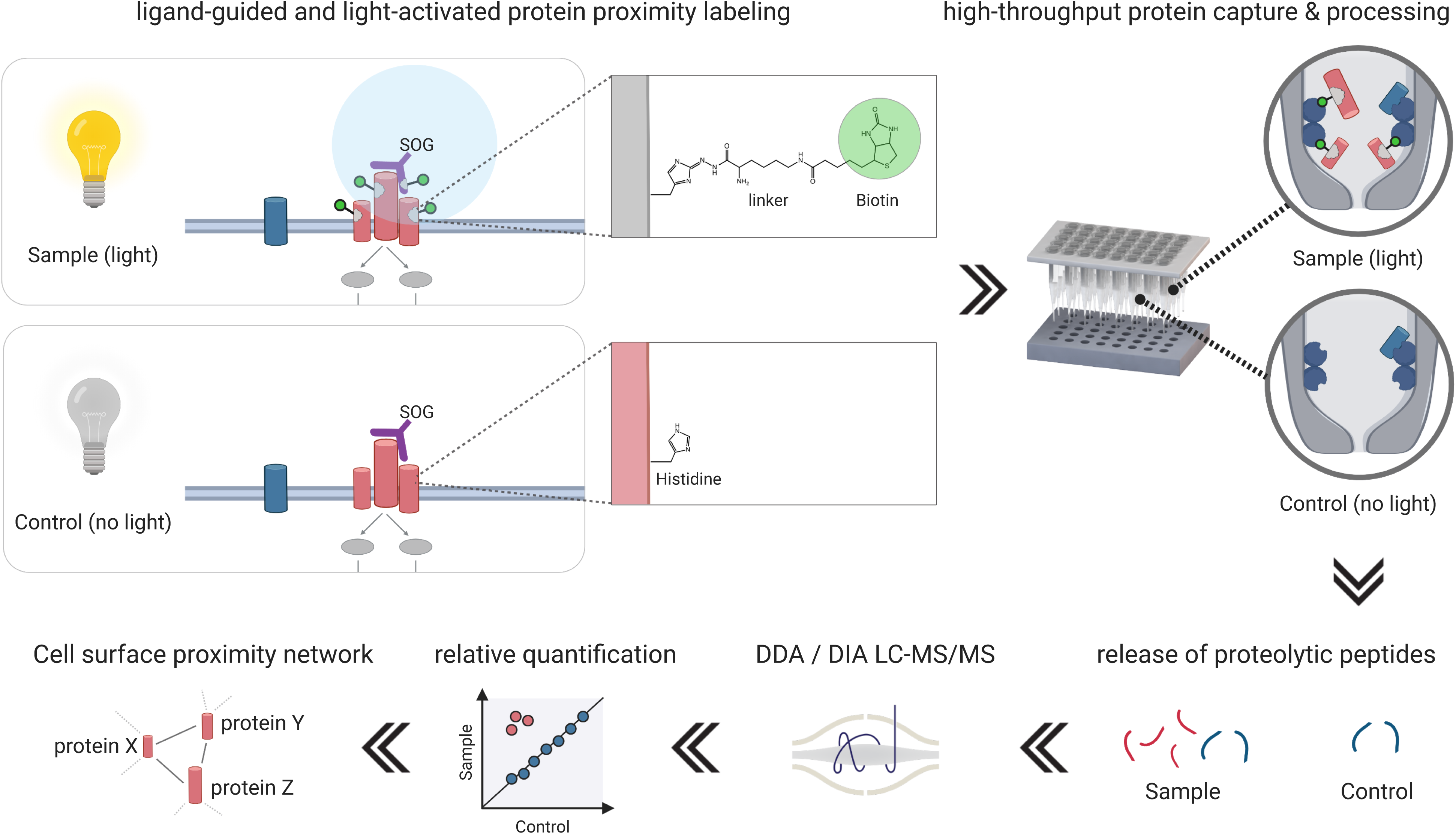
The LUX-MS strategy for light-activated identification of surfaceome signaling domains. Illumination of antibody- or ligand-coupled singlet oxygen generators (SOG) promotes spatially restricted oxidation of targeted surfaceome landscapes with spatiotemporal specificity. The light-tunable activation of amino acids such as Histidine (His) thereby enables chemical protein labeling under physiological conditions using biotin-functionalized hydrazide linkers. Labeled proteins are automatically captured and processed using a liquid-handling robotic platform ensuring fast and reproducible mass spectrometry-based quantification across a large number of samples. Relative quantification between illuminated and non-illuminated samples ultimately enables proteome-wide discovery of acute antibody- and ligand-targeted surfaceome signaling domains on living cells.

### LUX-MS enables proteome-wide mapping of antibody binding targets and surfaceome nanoscale organisation

Surfaceome interactions determine antibody specificity and signaling capacity and their elucidation is critical for therapeutic antibody development. In context of the highly successful therapeutic antibody rituximab^41^, decades of intensive research revealed the cell surface organisation of its primary receptor CD20 on targeted human B-lymphoma cells. To evaluate LUX-MS to provide further insights into rituximab action, we coupled anti-CD20 antibodies to the small molecule SOG thiorhodamine (**Fig. 2a**) and incubated resulting Ab-SOG constructs with human peripheral blood mononuclear cells (PBMCs) that were isolated from healthy donors. Flow cytometric analysis revealed antibody binding to 8% of all detected cells (**Fig. 2b**) corresponding to the typical fraction of CD20-positive cells in human PBMCs (5-10%)^42^. Subsequent illumination for 5 min in the presence of BH followed by flow cytometry demonstrated substantial cell surface biotinylation of a majority (96 %) of Ab-SOG bound but not unbound cells indicating efficient proximity labeling on living cells in complex biological environments with spatial specificity. We then examined LUX-labeling dynamics using Ab-SOG on a human B-lymphoma cell line and observed light-dependent cell surface biotinylation to be fine-tunable by the duration of illumination (**Fig. 2c** and **Supplementary Fig. 3**). Buffer conditions are known to modulate production and lifetime of singlet oxygen species^43^. Indeed, we achieved 5-fold enhanced photo-oxidation efficiency by replacing H_2_O with D_2_O in the buffer system and two-fold increased cell surface biotinylation when using 2-(aminomethyl)imidazole as a catalyst to boost hydrazone formation instead of the previously applied 5-methoxyanthranilic acid^19,43,44^ (**Supplementary Fig. 4**). We then performed a CD20-targeted LUX-MS experiment by treating cultured human B-lymphoma cells with Ab-SOG prior illumination for 0-5 min. In total, 1674 proteins were quantified by LUX-MS with at least two peptides per protein (**Supplementary Table 1**). While most proteins were uniformly abundant in the non-illuminated conditions, 63 proteins including 55 (87%) surface proteins showed striking enrichment that correlated with the duration of illumination and that culminated in a 5-fold abundance increase after 5 min of light (**Fig. 2d** and **Supplementary Fig. 5**). LUX-MS identification was thereby restricted to 43 of the 215 CSPA-reported surfaceome members and included non-glycosylated surface proteins that typically evade CSC- or LRC-based surfaceome interrogations (**Supplementary Fig. 6**). Proximity detection by LUX-MS therefore enables spatial proteotyping with sub-surfaceome resolution. In a second experiment using replicates, we used statistical testing to identify bona-fide SOG-proximal candidates (enrichment fold-change > 1.5 and p-value < 0.05, **Fig. 2e, Supplementary Table 1** and **Supplementary Data 1**). Among the top hits, we found immunoglobulin G2 (IgG2) isoforms of the Ab-SOG, the primary binding target CD20, human leukocyte antigen class II, tetraspanin CD37, CD298 and CD71 that were all shown to physically or functionally associate with

**Fig. 2.**
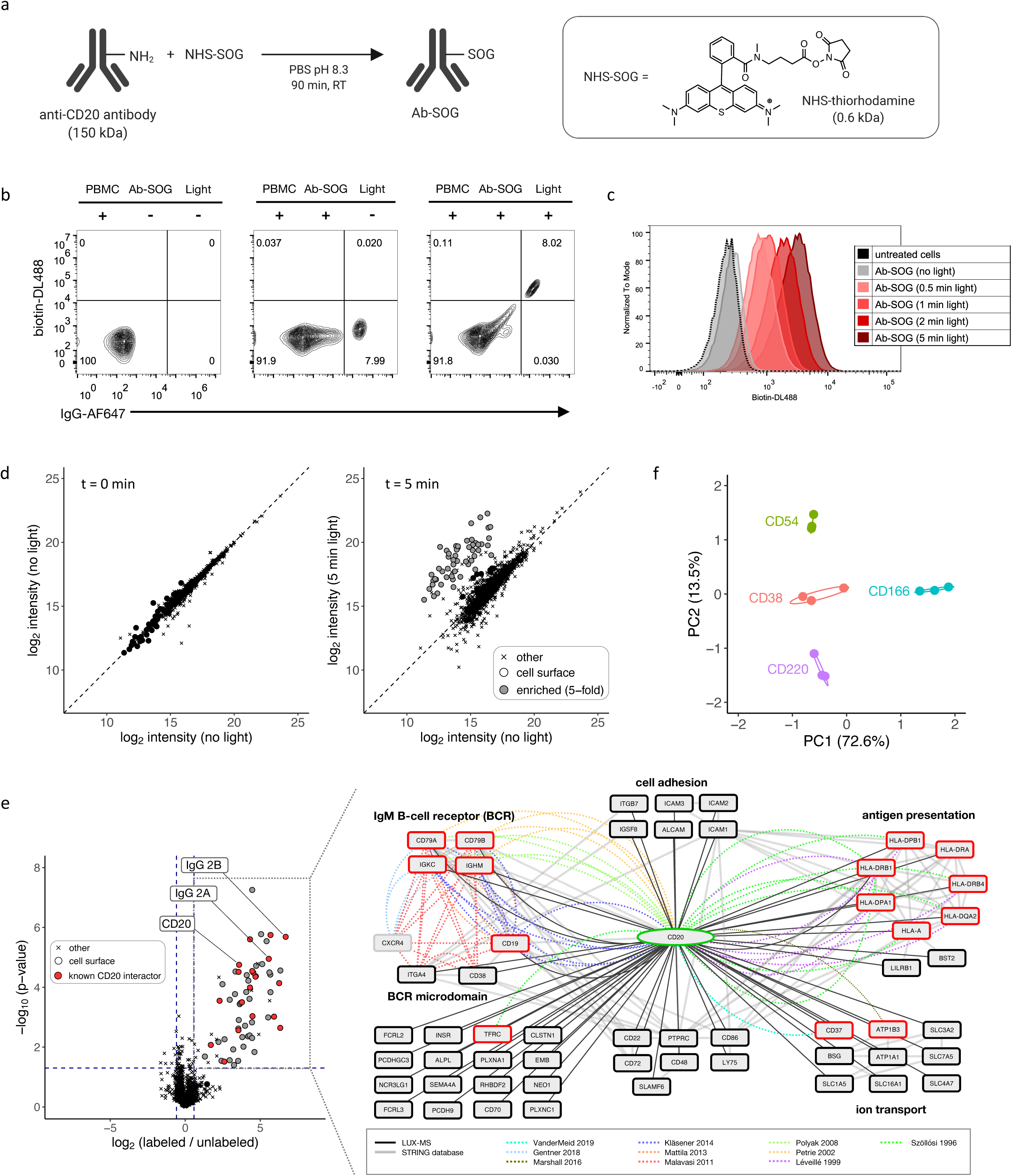
Proteome-wide discovery of antibody binding targets and surfaceome nanoscale organization. **a**, Schematic of decorating an antibody (Ab) with the singlet oxygen generator (SOG) thiorhodamine. **b**, Flow cytometric analysis of human peripheral blood mononuclear cells (PBMC) stained for cell surface biotin and immunoglobulin before and after LUX-labeling with anti-CD20 Ab-SOG. **c**, Histogram plot showing light-dependent cell surface biotinylation of LUX-labeled B-lymphoma SUDHL6 cells. **d**, Scatter plot showing light-dependent abundance of LUX-MS quantified proteins from Ab-SOG treated B-lymphoma SUDHL6 cells. Dots and crosses represent cell surface and otherwise annotated proteins, respectively. The color grey indicates 5-fold quantitative enrichment. **e**, left, Volcano plot showing relative abundance of LUX-MS quantified proteins from anti-CD20 Ab-SOG treated B-lymphoma SUDHL6 cells with and without illumination for 5 min. Dots and crosses represent cell surface and otherwise annotated proteins, respectively. Red and grey dots represent known and previously unknown CD20 associated proteins, respectively. The primary target CD20 and chains of the used antibody are highlighted. Right, literature and LUX-MS based cell surface interaction network of CD20. **f**, PCA analysis of CD38, CD54, CD166 and CD220 surfaceome neighborhoods on living B-lymphoma SUDHL6 cells identified by antibody-guided LUX-MS. A list of MS-quantified proteins (**Supplementary Table 1**) and a fully interactive volcano plot (**Supplementary Data 1**) are provided to facilitate data exploration.

CD20 on the surface of B-lymphoma cells^45–53^. Furthermore, LUX-MS revealed CD20 proximity to the IgM B cell receptor (BCR) complex (CD79A, CD79B, IgM heavy and light chain) including BCR-associated CD19, CD38 and ITGA4 supporting a model where rituximab binding to CD20 is able to eradicate B-lymphoma cells by altering survival-promoting BCR signaling^54,55^. Performing high-throughput LUX-MS on the same B-lymphoma cell line using a variety of SOG-coupled antibodies (against CD38, CD54, CD166, and CD220) thereby revealed receptor-specific surfaceome neighborhoods that covered distinct protein functionalities (**Fig. 2f** and **Supplementary Table 1**). Taken together, the spatiotemporal specificity of LUX-MS proximity detection enables the proteome-wide elucidation of antibody-surfaceome interactions and in a discovery-driven fashion and provides the throughput required for systematic mapping of the surfaceome nanoscale organization.

### LUX-MS decodes surfaceome signaling domains of small molecule drugs and biomolecules

Receptor interactions of small molecule drugs are crucial elements in many therapeutic strategies and receptor proximities on the surface of cells have recently become a target for precision therapies by drug-conjugated bispecific antibody (DBA)^56^ and extracellular-drug-conjugate (EDC) technologies^47^. EDC thereby exploit cell surface proximities between disease-relevant antibody targets and small molecule payload receptors for efficient cell ablation. However, the identity of small molecule receptors and their organisation within the surfaceome of live cells remains limited as enzyme-based proximity detection strategies are not attuned to using small-molecule ligands. To test whether LUX-MS can reveal small-molecule-targeted surfaceome domains, the SOG thiorhodamine was covalently conjugated to CG1, a small molecule drug that selectively inhibits the active ion pumping subunit (ATP1A1) of the ubiquitously expressed human Na^+^/K^+^-ATPase complex (NKA) (**Fig. 3a**)^47^. Cultured promyelocytic leukemia cells were incubated with increasing concentrations of the conjugate and LUX-labeling was performed as described above. FACS analysis showed light-induced and concentration-dependent cell-surface biotinylation that was saturated at 5 μM conjugate demonstrating binding to and labeling of surface molecules (**Fig. 3b**). Labeled cells were then subjected to the LUX-MS workflow resulting in significant enrichment of 48 surface-localized proteins upon illumination (**Fig. 3c, Supplementary Table 2** and **Supplementary Data 2**) including the beta3 subunit of the NKA (ATP1B3), indicating that LUX-MS enables discovery-driven detection of small-molecule-receptor interactions. Furthermore, we found the cell-surface receptor Basigin (CD147), which is a functional target of CG1-loaded extracellular-drug conjugates on the cell type used for this experiment^47^. Therefore the data suggests that small-molecule-targeted receptors and associated surfaceome signaling domains can be revealed by LUX-MS eventually enabling target discovery for DBA or EDC development.

**Fig. 3.**
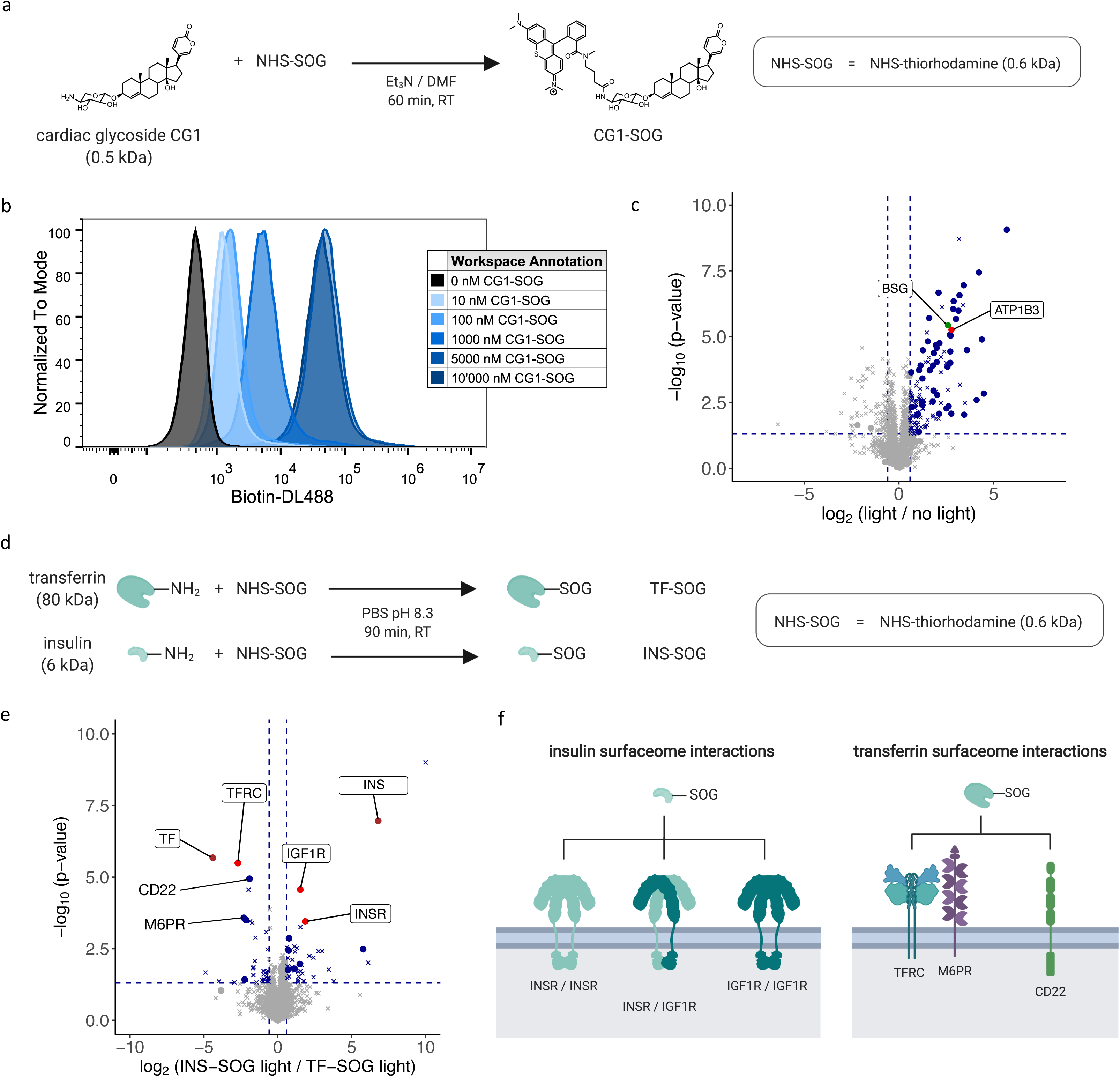
Decoding surfaceome signaling domains of small molecule drugs and biomolecules. **a**, Schematic of coupling the small molecule drug cardiac glycoside CG1 to the singlet oxygen generator (SOG) thiorhodamine. **b**, Histogram plot showing light-dependent cell surface biotinylation of LUX-labeled promyelocytic leukemia HL60 cells. **c**, Volcano plot showing relative abundance of LUX-MS quantified proteins from CG1-SOG treated promyelocytic leukemia HL60 cells with and without illumination for 5 min. Dots and crosses represent cell surface and otherwise annotated proteins, respectively. Red, green and blue dots represent the binding target of CG1, known and previously unknown surfaceome interactors, respectively. The former two are highlighted. **d**, Schematic of coupling the biomolecules insulin and transferrin to the singlet oxygen generator (SOG) thiorhodamine. **e**, Volcano plot showing relative abundance of LUX-MS quantified proteins from insulin-SOG and transferrin-SOG treated B-lymphoma SUDHL6 cells illuminated for 5 min. Dots and crosses represent cell surface and otherwise annotated proteins, respectively. Brown, red and blue dots represent the ligand, primary binding target and potential surfaceome interactors, respectively. The former two are highlighted together with known surfaceome interactors. **f**, Schematic representation of surfaceome interaction network identified by LUX-MS. A list of MS-quantified proteins (**Supplementary Table 2**) and fully interactive volcano plots (**Supplementary Data 2 and 3**) are provided to facilitate data exploration.

We next asked whether LUX-MS can be applied to uncover surfaceome interactions of more complex ligands i.e. secreted biomolecules that are key elements of intercellular communication. We therefore coupled thiorhodamine SOG to the peptide hormone insulin and the plasma glycoprotein transferrin that regulate cellular homeostasis through interactions with insulin receptor (INR) or insulin-like growth factor 1 receptor (IGF1R) homo- or heterodimers, and the transferrin receptor (TFRC), respectively (**Fig. 3d**). LUX-MS was performed on human B-lymphoma cells that were treated with either conjugate and illuminated for 5 min. Results demonstrated specific enrichment of insulin or transferrin in conjunction with their cognate receptors INSR and IGF1R or TFRC, respectively (**Fig. 3e, Supplementary Table 2** and **Supplementary Data 3**). Transferrin-guided LUX-MS further revealed CD22, a B-cell specific lectin shown to weakly bind to sialylated N-glycoproteins including transferrin^57^, and the cation-independent mannose-6-phosphate receptor M6PR, a sorting enzyme involved in the clathrin-mediated endocytosis and recycling of transferrin-bound receptors that was shown to co-localize with TFRC prior to internalization into sorting endosomes^58^. Thus, in addition to small molecules, LUX-MS can be used to unravel surfaceome signaling domains of biomolecules providing molecular insights into long-range intercellular communication events.

### Surfaceome signaling interactions can be detected in eukaryotic and prokaryotic systems

Protein proximity labeling and detection by LUX-MS primarily relies on light-induced modifications of histidine and tryptophan, whose cellular biosynthesis is evolutionary well conserved^59,60^. Thus, LUX-MS may uncover ligand-surfaceome signaling interactions beyond eukaryotic systems such as bacteria where LRC-like approaches are not applicable, solving a critical step in the development of anti-infective measures against the growing threat of drug-resistant bacteria. The naturally occurring peptide Thanatin was recently shown to exert antimicrobial activity by binding to LptA and eventually blocking the outer membrane biogenesis complex Lpt in *Escherichia coli* (*E.coli*)^61^. To test the capability of LUX-MS to identify critical surfaceome interactions in bacteria, we coupled Thanatin to the singlet oxygen generator methylene blue (**Fig. 4a**) and added it to *E. coli cells* with or without competition by excess of unconjugated Thanatin. All cells were subsequently illuminated for 5 min with red light at 656 nm and subjected to LUX-MS for identification of proximity labeled proteins. Numerous outer membrane and periplasmic proteins were significantly enriched (**Fig. 4b, Supplementary Table 3** and **Supplementary Data 4**), indicating membrane-localized labeling by SOG-coupled Thanatin. LUX-MS thereby revealed the functional binding partners of Thanatin LptA and LptD as top enriched hits and suggested proximity to outer membrane proteins including BamA and BamC (**Fig. 4b**). Indeed, the Bam complex was shown to catalyze the folding and insertion of outer membrane proteins including LptD forming supramolecular surfaceome clusters on living bacteria ^62^. LUX-MS therefore enables surfaceome target deconvolution and maps biomolecular signaling interactions across the domains of life.

**Fig. 4.**
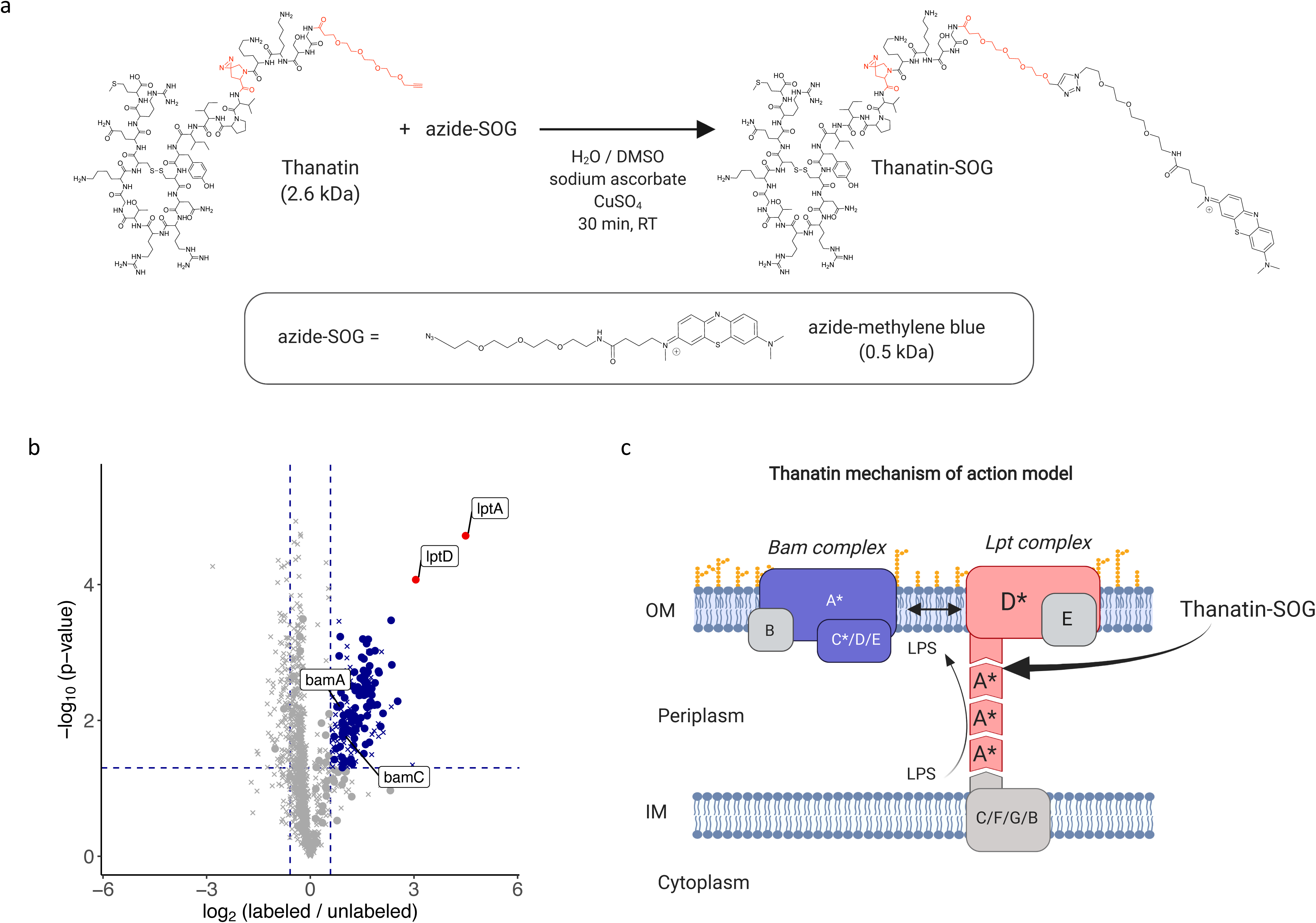
Hypothesis-free identification of surfaceome signaling interactions in prokaryotic systems. **a**, Schematic of coupling the peptidomimetic antibiotic Thanatin to the singlet oxygen generator (SOG) thiorhodamine. **b**, Volcano plot showing relative abundance of LUX-MS quantified proteins from Thanatin-SOG treated *Escherichia coli* illuminated for 15 min with and without Thanatin competition. Blue dots represent significantly enriched proteins and red dots represent the direct binding targets of Thanatin previously found by photo-crosslinking experiments. The main targets and selected surfaceome interactors are highlighted. **c**, schematic representation outlining the molecular mechanism of action of Thanatin in *E.coli* cells with spatial context. A list of MS-quantified proteins (**Supplementary Table 3**) and a fully interactive volcano plot (**Supplementary Data 4**) are provided to facilitate data exploration.

### Unravelling the surfaceome interaction network of intact viral particles using LUX-MS

Extracellular host-pathogen interactions are key signaling events that mediate cellular tropism and infection and, as such, are primary research targets for classification and diagnosis of infectious diseases and vaccine discovery. We therefore coupled thiorhodamine SOG to bacteriophages that recognize the ubiquitous cell-wall component teichoic acid of the Gram-positive bacteria *Listeria monocytogenes* (*Lm*), a highly infectious food-borne pathogen that causes listeriosis^63^ (**Fig. 5a**). *Lm* were incubated with SOG-coupled phages at multiplicity of infection (MOI) of 1 or 10 and LUX-labeled by illumination for 0 to 15 min. Flow cytometry analysis revealed the light-dependent cell-surface biotinylation of individual bacteria that reached saturation after 15 min of illumination at an MOI of 10 (**Fig. 5b**). Subsequent LUX-MS analysis revealed light-induced, significant enrichment of 146 proteins including 19 phage and 127 *Lm* proteins (**Fig. 5c, Supplementary Table 4** and **Supplementary Data 5**). We found components of bacteriophage capsid (Cps, Gp5, Gp9), sheath (Gp3), baseplate (Gp20), and tail (Gp17, Gp18, Gp19, Gp23, Tmp, Tsh) and enzymes involved in DNA binding (Gp37), replication (Gp45), recombination (Gp44), transcription (Gp7), and immunoprotective functions (Gp32) indicating thorough LUX-labeling of host-cell attached bacteriophages. For *Lm* proteins, gene ontology analysis revealed 81 extracellular and only 8 intracellular proteins (**Fig. 5d**) including major virulence factors such as cell wall-associated internalins A and B that mediate host cell invasion, membrane-localized ActA that governs cell-to-cell spreading and ATP-binding cassette (ABC) transporters that mediate multi-drug resistance ^64^ (**Fig. 5e**). Using HA-tagging and immunofluorescence microscopy, we further validated the *Lm* surface expression of a putative cell wall protein and ABC transporter that were strongly enriched by LUX-MS (**Fig. 5f**). Altogether, our results demonstrate the capability of LUX-MS to unravel host surfaceome interactions of intact viruses on a proteome-wide scale enabling the study of extracellular host-pathogen interactions eventually facilitating the design of anti-infective strategies.

**Fig. 5.**
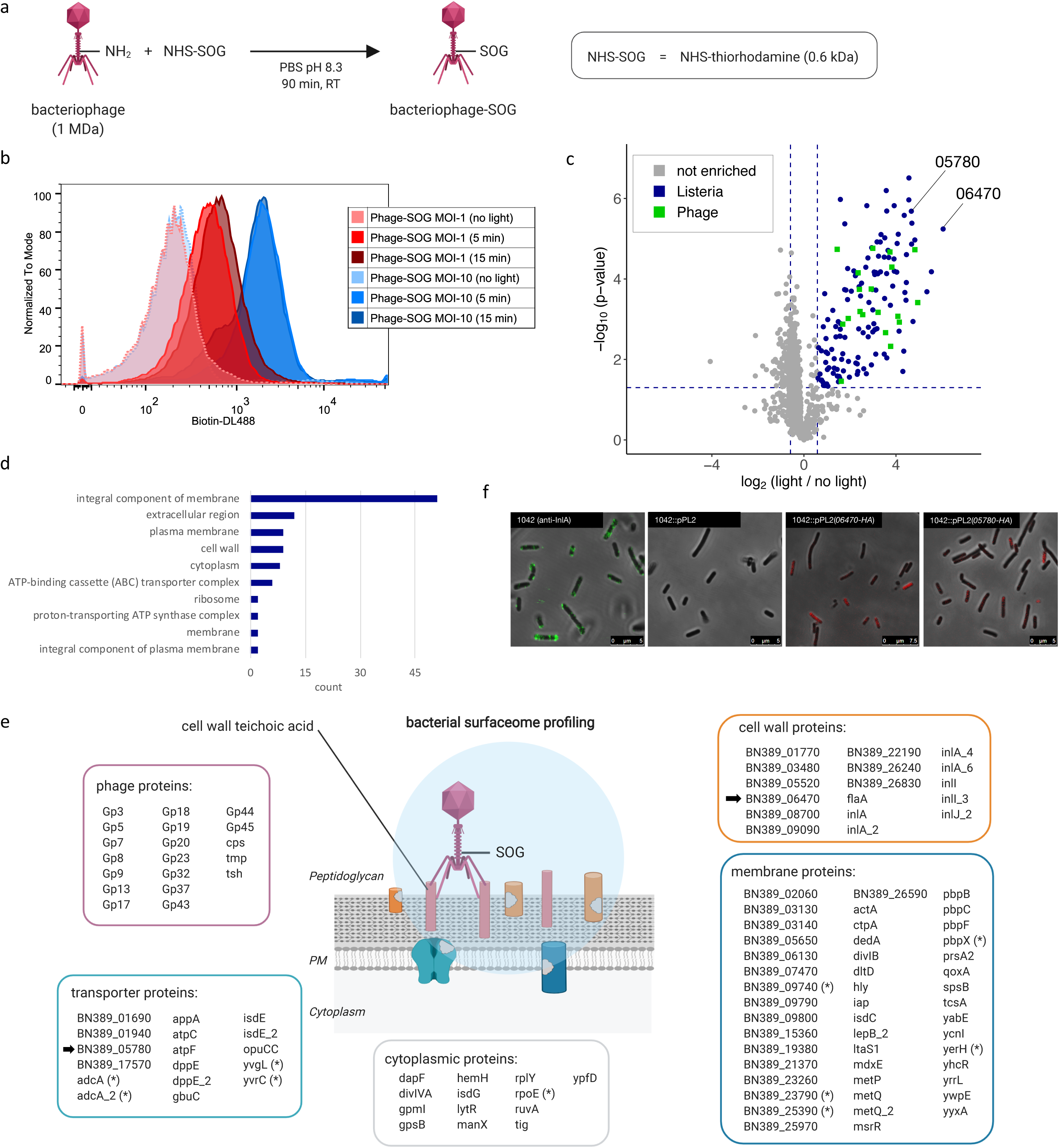
Mapping virus-targeted surfaceomes on living hosts using bacteriophage-guided LUX-MS. **a**, Schematic of coupling the *Listeria monocytogenes* (*Lm*)-specific bacteriophage to the singlet oxygen generator (SOG) thiorhodamine. **b**, Histogram plot showing light-dependent cell surface biotinylation of *Lm* cells after LUX-labeling with bacteriophage-SOG. **c**, Volcano plot showing relative abundance of LUX-MS quantified proteins from bacteriophage-SOG treated *Lm* with and without illumination for 15 min. Blue and green dots represent significantly enriched *Lm*- or phage-derived proteins, respectively, while grey dots indicate non-enriched proteins. Candidates selected for microscopic characterization are highlighted. **d**, GO-term enrichment analysis of significantly enriched *Lm* proteins. **e**, Schematic representation of the spatial coverage of LUX-MS based on subcellular annotations of significantly enriched proteins. Candidates selected for microscopy are pointed by arrows. **f**, Representative confocal microscopy images of *Lm* (1042) left untreated and stained for cell surface Internalin-A (green), or transfected with plasmid containing no (pPL2) or HA-tagged candidate proteins and stained for cell surface HA (Red). A list of MS-quantified proteins (**Supplementary Table 4**) and a fully interactive volcano plot (**Supplementary Data 5**) are provided to facilitate data exploration.

### Elucidating intercellular surfaceome signaling domains in functional immunosynapses

Contact-dependent intercellular communication is an important regulator of immune system function and relies on highly complex intercellular surfaceome signaling architectures - in case of T lymphocyte activation by antigen presenting cells (APCs) - the “immunological synapse”^6^. The contextual interplay of immunosynaptic receptor interactions between APCs and CD8^+^ T cells leads to T cell activation, proliferation and differentiation enabling these cells to exert effector functions against infected, tumorigenic or foreign cells^65^. Thus, the ability to resolve functional immunosynapses at molecular level with spatiotemporal specificity would enable discoveries in the field of immunology and immuno-oncology. We thus coupled thiorhodamine SOG to the lymphocytic choriomeningitis virus-(LCMV)derived peptide gp33-41 (gp33, sequence: KAVYNFATM) (**Fig. 6a**), an immunodominant epitope of the LCMV glycopotein (GP) that is presented by MHC class I H-2D^b^ and H-2K^b 66^ and that is recognized by T cell receptor (TCR) transgenic CD8^+^ T cells of P14 mice^67^. In a two-cell system, we showed that SOG-coupled gp33 is still presented by mouse dendritic cells and is able to comparably activate freshly isolated P14 CD8^+^ T-cells than its unconjugated counterpart (**Supplementary Fig. 8**). Applying light led to cell surface biotinylation of both cell types, validating the photocatalytic function of SOG-coupled gp33 and our ability to efficiently perform trans-cellular surfaceome labeling during immunosynaptic cell-to-cell contacts (**Supplementary Fig. 9**). We then implemented stable isotope labeling by amino acids in cell culture (SILAC) for isotopic barcoding of interacting dendritic cells (DC) and T cells (TC) and performed LUX-MS in data-independent acquisition (DIA) mode after 30 min of co-culture to analyse mature immunological synapses (**Fig. 6b**). Overall, we identified 4811 protein groups that could clearly be assigned to TC (e.g. TCR, CD3, and CD8), DC (e.g. MHC class I H-2D^b^ and H-2K^b^), or both based on observed SILAC-ratios (**Fig. 6c** and **Supplementary Table 5**). We observed light-dependent enrichment of numerous cell surface proteins including components of the primary signaling axis such as MHC class I on DC and the gp33-specific TCR as well as CD8 on TC (**Fig. 6d-e, Supplementary Table 5, Supplementary Data 6 and 7**). Furthermore, we found co-stimulatory signaling components essential for early T cell priming such as CD86 of the CD86-CD28 axis and the entire CD2-CD48 axis with reported cellular directionality^6^. Our results further indicate immunosynaptic localization of CD5 and CD6 on TC. Indeed CD5 was shown to interact with TCR, CD8, and CD2 in the immune synapse to fine-tune TCR-mediated signaling^68^, while the structurally related CD6 physically associates with CD5 and its ligand CD166 (also known as ALCAM)^69^ that was also captured on DC. LUX-MS further revealed immunosynaptic adhesion complexes including LFA-1 (TC), ICAM-1 (DC) and VLA-4 (both) that enhance intercellular proximity and T cell activation^65^ and identified highly membrane-embedded tetraspanins CD9, CD81, CD82, and CD151 on DC that organize immunosynaptic signaling domains on antigen-presenting cells^70^. In contrast, the ubiquitous protein tyrosine phosphatase receptor type C (PTPRC, CD45) was not identified on TC in line with its immunosynaptic exclusion to permit TCR phosphorylation and signaling^65^ (**Fig. 6e**). Taken together, our results demonstrate the utility of LUX-MS to comprehensively resolve the intercellular surfaceome signaling architectures of functional immune synapses with cell-type specificity and spatiotemporal control by light, providing a molecular framework for drug discovery in immunology and immuno-oncology.

**Fig. 6.**
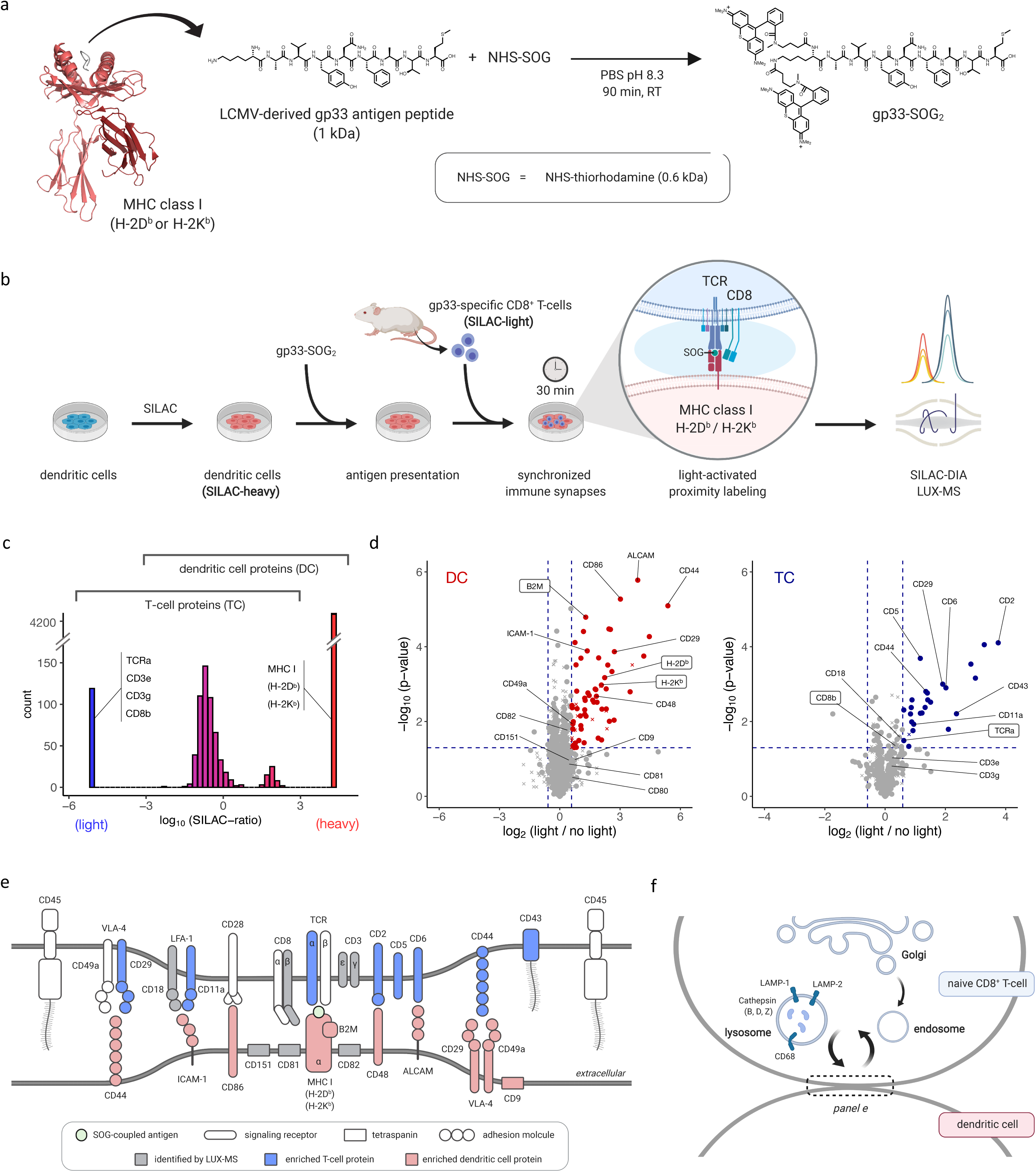
Elucidating intercellular surfaceome signaling domains in functional immunosynapses. **a**, chematic of coupling immunogenic peptide gp33 to the singlet oxygen generator (SOG) thiorhodamine. The crystal structure of gp33 (dark grey) presented by MHC class I H-2D^B^ alpha chain (red) in complex with beta-2 microglobulin (dark red) is shown (PDB identifier: 1FG2) **b**, Schematic of the SILAC-DIA based LUX-MS workflow. Isotopically labeled (heavy) mouse dendritic cells presenting gp33-SOG2 establish synchronized immunosynapses with isolated (light), gp33-specific CD8^+^ T cells enabling light-activated *in-situ* labeling and molecular analysis of intercellular surfaceome signaling interactions within synapses using data-independent acquisition (DIA) mass spectrometry. **c**, Histogram plot showing distribution of heavy-to-light abundance ratios of LUX-MS identified proteins. Ratio-boundaries for cell type assignment and representative proteins are shown. **d**, Volcano plot showing relative abundance of LUX-MS quantified proteins from the two-cell systems with and without illumination for 15 min. Dots and crosses represent cell surface and otherwise annotated proteins, respectively. Red and blue dots represent dendritic cell (DC) and T cell (TC) derived proteins, respectively. Known immunosynaptic constituents are highlighted with direct gp33 interactors shown in boxes. **e**, Schematic of the LUX-MS identified immunosynaptic surfaceome interaction network. **f**, Schematic highlighting the interconnectivity of the immunological synapse with endolysosomal pathways in T-cells and showing proteins reported at the cell-to-cell interface by LUX-MS. A list of MS-quantified proteins (**Supplementary Table 5**) and a fully interactive volcano plot (**Supplementary Data 6 and 7**) are provided to facilitate data exploration.

## Discussion

The nanoscale organisation and extracellular interaction network of the cellular surfaceome is a key signaling mediator in drug action, viral infection and intercellular communication. In this study, we developed and applied a SOG-based and light-controlled proximity detection technology termed LUX-MS that enables to elucidate the surfaceome nanoscale organization and its ligand-targeted surfaceome signaling doamins on living cells. In contrast to larger enzymes (APEX2: 28 kDa; miniTurbo: 27 kDa)^71^, the use of small molecular SOG (< 1 kDa) as labeling probes offers the following advantages: i) A broad ligand-compatibility enabling target deconvolution and surfaceome interaction profiling of ligands of basically any description ranging from small molecules, antibodies and viral particles to whole cells. ii) Covalent SOG-coupling with ligand-compatible chemistry preserving structure-function relationships and circumventing the need for genetic manipulation. iii) Light-controlled singlet oxygen generation allowing for fine-tunable *in-situ* labeling with tight spatiotemporal control. iv) Protein labeling via conserved amino acid residues enabling proteome-wide surfaceome interrogations in a broad range of evolutionary distinct organisms. Recently, photocatalysts producing short-lived carbene with nanosecond half-lives were utilized for antibody-guided mapping of protein microdomains that could not be resolved using enzyme-based strategies (intermediates with half-lives of 0.1 ms to 60 s)^27^. Proximity labeling by LUX-MS is mediated by photosensitized singlet oxygen species with a lifetime of 25 - 36 μs^43^ indicating spatial resolution in between enzyme- and carbene-based approaches. Still, the direct coupling of SOG to signaling competent ligands confines labeling to signaling relevant surfaceome domains and circumvents the use of antibodies that entail structural flexibility. Thus, LUX-MS complements currently available technologies and provides an analytical stepping stone for system-wide elucidation of the cellular surfaceome landscape.

Using antibodies to guide proximity labeling, we demonstrated the ability of LUX-MS to specifically reveal distinct surfaceome proximities on living cells that form a molecular framework for the reconstruction of the surfaceome nanoscale organisation. Given the broad availability of high-quality antibodies for established surface targets and the high-throughput capability of the LUX-MS workflow, profound surfaceome interaction mapping is feasible providing the cell surface level information that is currently lacking in ongoing systems biology efforts to map and understand the structure and function of the human interactome^72–75^.

By applying LUX-MS to small molecules, we mapped the surfaceome signaling domains of the small molecule drug cardiac glycoside CG1. In contrast to ligand-receptor capture methodologies, LUX-MS not only enabled proteome-wide target deconvolution of small molecules but also provided cell-type specific surfaceome proximity information that could be leveraged for targeted drug delivery e.g. using EDC ^47^. While this approach is expected to be transferable to other drug modalities, the accumulating knowledge on drug-targeted cellular surfaceomes eventually allows for the identification of disease-associated surfaceome proximites enabling a next generation precision medicine that goes well beyond the targeting of individual receptors.

Using LUX-MS, we decoded viral surfaceome interactions in bacterial hosts that are not detectable using state-of-the-art technologies such as CSC, LRC, and HATRIC due to the structural complexity of the cell envelope and a general lack of protein glycosylation. Using bacteriophages that recognize a ubiquitous cell wall component of *Listeria monocytogenes*, we mapped the bacterial surfaceome (>80 proteins) with substantial coverage and specificity that superseded common profiling strategies such as trypsin shaving (11 proteins identified), surface-restricted biotinylation (27 proteins identified), and cell fractionation (21 proteins identified)^76^. Depending on the binding specificity of the employed viral particle, LUX-MS enables the spatiotemporal elucidation of viral attachment sites, target receptors, and global surfaceomes of live prokaryotic and eukaryotic cells in complex biological systems. This is of particular interest considering the broad application of advanced phage display assays^77^ and related screening platforms^5,78^ in basic and translational research.

Finally, we employed LUX-MS for the spatiotemporal analysis of intercellular surfaceome signaling domains within immunological synapses. Previous attempts were limited to molecular analysis in model systems that use soluble, plate-bound, or membrane-associated factors to mimic immunosynaptic T-cell activation^65^ or to fluorescence-based reporting of transient intercellular immune cell interactions^27,79^. We here performed a gp33 immunogen-guided LUX-MS in an isotopically barcoded two-cell system of millions of synchronized and functional immunosynapses and thereby obtained a holistic view on the mature immunosynaptic surfaceome signaling architecture, fostering the formulation of novel biological hypotheses. For example, we found several lysosomal proteins (LAMP-1, LAMP-2, CD68, and cathepsins B, D, and Z) within the immunosynaptic cleft on naive CD8^+^ T-cells but not antigen-presenting dendritic cells (**Supplementary Table 5**). This indicates local fusion of lysosomal granules in T cells that is typically observed in cytotoxic T-cells during degranulation-mediated target cell killing^80^. However, while being a feature of antigen-experienced cells, degranulation and thus cytolytic activity was shown to be absent in naive CD8^+^ T cells. Indeed, we did not identify any cytolytic effector molecules such as perforin or granzymes, indicating the secretion of non-lytic granules in naive CD8^+^ T cells within 30 min of initial contact with antigen-presenting cells (**Fig. 6f**). Albeit, the cellular machinery for immunosynapse-directed transport of vesicles is in place within 6 min of initial cell contact, the exact mechanism of such non-lytic granule secretion and its functional role in antigen-dependent T-cell priming remains to be further explored.

In conclusion, we provide a high-throughput, optoproteomic technology termed LUX-MS to discover protein networks within surfaceome signaling domains on living cells and we demonstrate its utility and versatility by unravelling surfaceome nanoscale organisation and signaling interactions underlying drug action, host-pathogen interactions and intercellular communication. Given the biological and therapeutic importance of the surfaceome signaling landscape, we envision LUX-MS to impact basic and applied research areas by unravelling the spatiotemporal interconnectivity of the cell surface signaling gateway in health and disease.

## Methods

### Reagents

All chemicals were purchased from Merck unless stated otherwise. Singlet oxygen generators in form of NHS-functionalized thiorhodamine (cat: AD-Thio12-41) and azide-functionalized methylene blue (cat: AD-MB2-31) were purchased from ATTO-TEC GmbH (Siegen, Germany). Biocytin-hydrazide was purchased from Pitsch Nucleic Acids AG (Stein am Rhein, Switzerland). The following antibodies were used for LUX-MS experiments: mouse IgG Isotype control antibody (Invitrogen, cat: 10400C), anti-human CD20 mouse IgG2 monoclonal antibody clone 2H7 (Invitrogen, cat: 14-0209-82), anti-human CD38 chimeric monoclonal antibody kindly provided by Centrose LLC, anti-human CD54 mouse IgG1 monoclonal antibody, clone HCD54 (BioLegend, cat: 322704), anti-human CD166 mouse IgG1 monoclonal antibody clone 3A6 (BioLegend, cat: 343902), anti-human CD220 mouse IgG2 monoclonal antibody, clone B6.220 (BioLegend, cat: 352602).

### Cell culture

All chemicals for cell culture were purchased from ThermoFisher Scientific unless stated otherwise. Cell lines were purchased from ATCC and grown at 37 °C and 5% ambient CO_2_. Patient-derived B-lymphoma cell line SU-DHL-6 (ATCC, CRL-2959) was grown in Roswell Park Memorial Institute (RPMI) 1640 medium with 1.5 mM GlutaMAX, 1% penicillin-streptomycin and 10% fetal bovine serum. Patient-derived promyelocytic leukemia cell line HL60 (ATCC CCL-240) was grown in Iscove’s Modified Dulbecco’s Medium (IMDM) with 1.5 mM GlutaMAX, 1% penicillin-streptomycin and 10% fetal bovine serum. Mouse dendritic cell line MutuDC1940^81^ was kindly provided by Hans Acha-Orbea (Department of Biochemistry, University of Lausanne, Switzerland) and was cultured and isotopically labeled in Iscove’s Modified Dulbecco’s Medium (IMDM) for SILAC (Thermo, cat: 88367) supplemented with glucose (final: 4.5 g/l), L-Arg-^13^C_6_-^15^N_4_ (final: 42 μg/ml), L-Lys-^13^C_6_-^15^N_2_ (final: 73 μg/ml), 1.5 mM GlutaMAX, 10 mM HEPES, 50 μM β-mercaptoethanol, 1% penicillin-streptomycin, non-essential amino acids and 10% dialyzed fetal bovine serum.

### Mice

Female or male mice of six to sixteen weeks of age were used for animal experiments described in this study. CD45.1 P14 mice with T cell receptors on CD8^+^ T cells specific for the glycoprotein GP33-41 epitope of lymphocytic choriomeningitis virus (LCMV) ^67^ and CD45.1 P14 crossed with Nur77-GFP mice^82^ were bred at the ETH Phenomics Center. All animals were bred and held under specific-pathogen-free conditions prior to use. All animal experiments were performed in accordance with institutional policies and Swiss federal regulations, following guidelines and being approved by the veterinary office of the Canton of Zürich (animal experimental permissions: 115/2017).

### Light-controlled production of singlet oxygen

The singlet oxygen generator (SOG) thiorhodamine was mixed with 1 μM Singlet Oxygen Sensor Green (Invitrogen, cat: S36002) in phosphate buffered saline at increasing D_2_O/H_2_O ratios and illuminated at a working distance of 30 cm with 590-nm Precision LED spotlights controlled by a BioLED Light Control Module (Mightex Systems) for 0 to 15 minutes. The fluorescence of photo-oxidized SOSG was monitored using a Synergy HT Multi-Mode Plate Reader (BioTek) at excitation 485 ± 20 nm and emission 528 ± 20 nm. Quantified intensities were blanked and normalized to non-illuminated control using a customized R-script.

### Discovery of light-induced protein modifications

Human holo-transferrin (Sigma, cat: T4132) was coupled with thiorhodamine SOG at a molar ligand:SOG ratio of 1:1 and purified using 7-kDa ZebaSpin Columns (Thermo, cat: 89882) according to manufacturer’s recommendations. SOG-coupled transferrin in D_2_O-based PBS containing 0 or 5 mM biocytin-hydrazide was either illuminated with 590-nm Precision LED spotlights operated via a BioLED light source control module (Mightex Systems, Pleasanton, USA) for 15 min or left in the dark. The buffer was complemented with 50 mM 2-(aminomethyl)imidazole dihydrochloride catalyst for 50 min. Treated proteins were digested overnight at 37 °C using sequencing-grade trypsin (Promega, cat: V511C) at an enzyme-to-protein ratio of 1:100. Peptides were C18-purified using 5–60 μg UltraMicroSpin Columns (The Nest Group, cat: SEMSS18V) according to manufacturer’s instructions and subjected for mass spectromic analysis using an Orbitrap Fusion Tribrid mass spectrometer (Thermo Scientific) in data-dependent acquisition (DDA) mode with a mass resolution of 120,000 and 30’000 on precursor and fragment level, respectively (see below for details). Acquired raw files of illuminated samples were subjected to an open modification search using MSfragger (v.20190628)^83^ and Crystal-C^84^ within the FragPipe pipeline (v.9.4) using standard settings and a SwissProt-reviewed human protein database (downloaded June 2014) containing common contaminants. Observed modifications were implemented in a subsequent closed search of all acquired samples using Comet (v.2015.01) within the Trans Proteomic Pipeline v.4.7 (SPC/ISB Seattle) as variable modification (carbonylation of His, Cys, and Trp; single oxidation of Met, Trp, and Tyr; double oxidation of Met, Trp, and His; triple oxidation of Met, Cys, and Trp) to identify fully-tryptic peptides with a maximum of two missed cleavage sites and a maximum of 5 variable modifications. The precursor and fragment mass tolerance was thereby set to 20 ppm and 0.02 Da, respectively. Peptides were quantified by integration of chromatographic traces using Progenesis QI v.4.0 (Nonlinear Dynamics) and filtered to a false discovery rate of < 1%. Transferrin peptides were assigned to dummy proteins representing unmodified or single modification types that were statistically tested for differential abundance using R package MSstats (v.3.8.6)^85^. Modification types with an abundance fold change > 1.5 upon illumination and an adjusted p-value < 0.05 were considered products of light-induced photo-oxidation. Analogously, modification types with a significantly reduced light-dependent production in the presence of biotin-hydrazide (p-value < 0.1) were considered hydrazide-reactive amino acid modifications.

### Collection and purification of human peripheral blood mononuclear cells (PBMCs)

Buffy coats were obtained from healthy donors provided by the Blood Transfusion Service Zurich. To isolate PBMCs, human buffy coat samples were diluted 1:1 in PBS (Gibco) and mononuclear cells were isolated with a Histopaque-1077 density gradient (Sigma-Aldrich) according to the manufacturer’s instructions. PBMCs at the interface were collected, washed once in PBS and resuspended in media until further processing.

### Antibody-guided surfaceome nanoscale mapping

Per sample, 10 μg antibody was coupled with thiorhodamine SOG at a molar antibody:SOG ratio of 1:5 and purified using 7-kDa ZebaSpin Columns (Thermo, cat: 89882) according to manufacturer’s recommendations. The SOG-coupled antibody constructs were kept at 4 °C and immediately used. Twenty millions cells were incubated with 10 μg SOG-coupled antibody for 30 min at 4 °C in the dark to minimize background light-induced oxidation. Cells were washed with ice-cold PBS, resuspended in chilled photo-oxidation buffer (5 mM biocytin-hydrazide in D_2_O-based PBS, pH 7.5) and illuminated at a working distance of 20 cm with 590-nm Precision LED spotlights or left in the dark. For the anti-CD20 time course LUX-MS, one and three biologically independent samples were analysed for each illumination time point and the non-illuminated control, respectively. For anti-CD20 LUX-MS two and three biologically independent samples were analysed for labeled and unlabeled conditions, respectively. For all other antibody-guided LUX-MS, three biologically independent samples were analysed for both labeled and unlabeled conditions. Cells were pelleted by centrifugation and resuspended in chilled labeling buffer (5 mM biocytin-hydrazide and 50 mM 2-(Aminomethyl)imidazole dihydrochloride in PBS, pH 6.0) for 50 min at 4 °C in the dark. For flow cytometric analysis of cell surface biotinylation, cells were extensively washed with PBS and stained with NeutrAvidin Protein DyLight 488 (Invitrogen, cat: 22832) 1:200 for 20 min prior analysis using an Accuri C6 Flow Cytometer (BD Biosciences) and FlowJo (v.10.07). The FSC/SSC pattern of the unlabeled condition was thereby used to gate for live cells. For LUX-MS analysis, cells were extensively washed with PBS, snap-frozen as cell pellets in liquid nitrogen and stored at −80 °C until further processing.

### Synthesis of singlet oxygen generator-coupled cardiac glycoside CG1

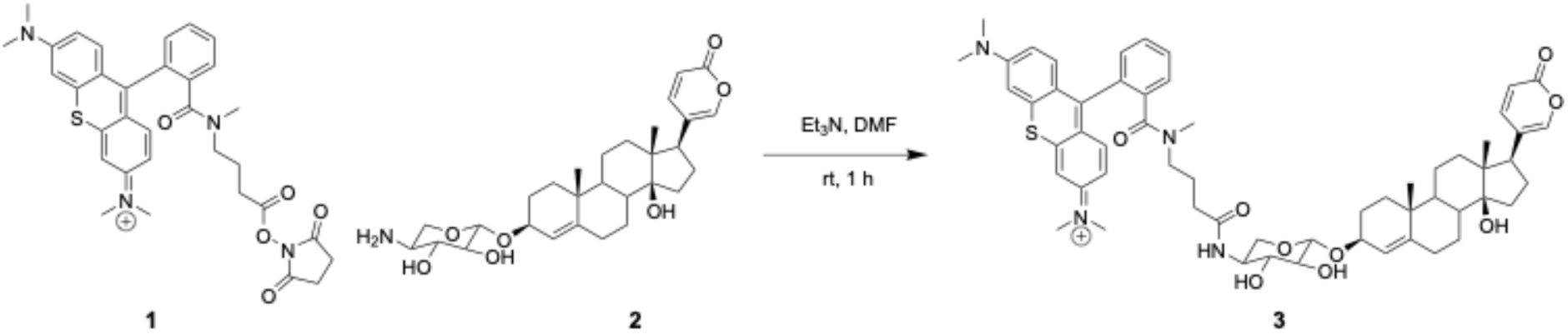

The primary amine containing cardiac glycoside CG1 was kindly provided by Centrose LLC (Madison, Wisconsin). To a solution of CG1 **2** (5 mg, 0.0081 mmol) and thiorhodamine-NHS ester **1** (5.7 mg, 0.0081 mmol) in N,N-dimethylformamide (DMF) (1 mL) was added Et_3_N (0.011 mL, 0.08 mmol). The reaction mixture was stirred at RT for 1 hour and product formation was confirmed by LC-MS. The final product CG1-SOG **3** was purified by preparative RP-HPLC on a Waters Eclipse XDB (C18, 250 x 21.2 mm, 7 μm) column with a gradient of 5 – 50% ACN/H_2_O + 0.1% TFA in 5 column volumes. The product was of >95% purity as judged by reversed phase UPLC and HR-MS (**Supplementary Fig. 10**). HR-ESI-MS: m/z (M+H^+^) 999.4938 (calc. mass = 999.4936).

### Identification of small molecule drug-targeted surfaceome structures

Fifty millions cells were incubated with 0 - 10 μM CG1-SOG (10 μM for LUX-MS experiment) for 30 min at 4 °C in the dark to minimize background light-induced oxidation. Cells were washed with ice-cold PBS, resuspended in chilled photo-oxidation buffer and illuminated at a working distance of 20 cm with 590-nm Precision LED spotlights or left in the dark (three biologically independent samples each). Cells were pelleted by centrifugation and resuspended in chilled labeling buffer for 50 min at 4 °C in the dark. For flow cytometric analysis of cell surface biotinylation, cells were extensively washed with PBS and stained with NeutrAvidin Protein DyLight 488 (Invitrogen, cat: 22832) 1:200 for 20 min prior analysis using an Accuri C6 Flow Cytometer (BD Biosciences) and FlowJo (v.10.07). The FSC/SSC pattern of the unlabeled condition was thereby used to gate for live cells. For LUX-MS analysis, cells were extensively washed with PBS, snap-frozen as cell pellets in liquid nitrogen and stored at −80 °C until further processing.

### Biomolecule-guided discovery of intercellular surfaceome signaling domains

The human peptide hormone insulin and glycoprotein holo-transferrin were coupled with thiorhodamine SOG at a molar ligand:SOG ratio of 1:1 and 1:13, respectively according to manufacturer’s recommendations. The SOG-coupled ligand constructs were kept at 4 °C and immediately used. Twenty millions cells were incubated with 1 μM Insulin-SOG or 75 nM Transferrin-SOG for 10 min at 4°C in the dark to minimize background light-induced oxidation. Cells were washed with ice-cold PBS, resuspended in chilled photo-oxidation buffer and illuminated at a working distance of 20 cm with 590-nm Precision LED spotlights for 5 min or left in the dark (three biologically independent samples each). Cells were pelleted by centrifugation and resuspended in chilled labeling buffer for 50 min at 4 °C in the dark. Cells were extensively washed with PBS, snap-frozen as cell pellets in liquid nitrogen and stored at −80 °C until further processing.

### Synthesis and characterization of singlet oxygen generator-coupled antimicrobial Thanatin

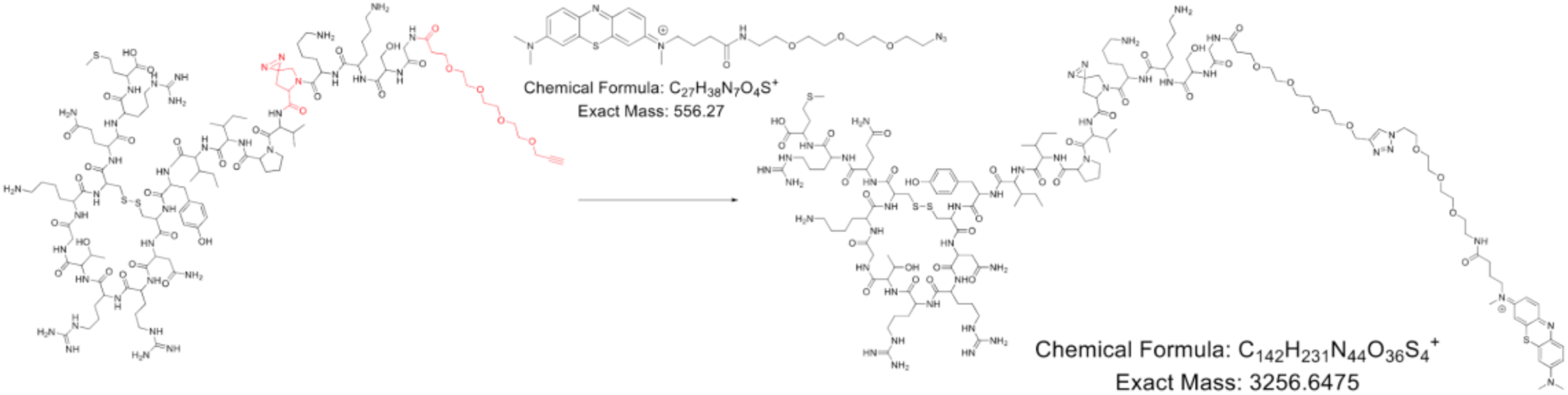

A Thanatin derivative was kindly provided by the John Robinson group (University of Zurich, Switzerland) and (6.0 mg, 0.0023 mmol) were dissolved in 500 μl H_2_O. Sodium ascorbate (0.1 M, 30 μl, 0.0022 mmol) and azide-functionalized methylene-blue as Singlet Oxygen Generator (0.5 mg, 0.0009 mmol) in 100 μl DMSO and CuSO_4_ (0.1 M, 30 μl, 0.003 mmol) was added. After 15 min the reaction mixture was directly injected into a prep HPLC C18 column with a gradient of 10-40% ACN/H_2_O (0.1% TFA). The purity of the Thanatin-SOG compound was confirmed by UPLC analysis and the identity was verified by HR-ESI with a calculated mass of 652.1353 (M+6H)^6+^ and a measured m/z value of 652.1360 (M+6H)^6+^ (**Supplementary Fig. 11**).

### Proteome-wide and light-activated target deconvolution in prokaryotic systems

*Escherichia coli* cells (Migula) Castellani and Chalmers (ATCC: 25922) were incubated with 3 μM Thanatin-SOG in the presence or absence of 30 μM unconjugated Thanatin derivative for 30 min at 37 °C in the dark to minimize background light-induced oxidation. Cells were washed with ice-cold PBS, resuspended in chilled photo-oxidation buffer and illuminated at a working distance of 20 cm with 659-nm Precision LED spotlights for 5 min or left in the dark (three biologically independent samples each). Cells were pelleted by centrifugation and resuspended in chilled labeling buffer for 50 min at 4 °C in the dark. Cells were extensively washed with PBS, snap-frozen as cell pellets in liquid nitrogen and stored at −80 °C until further processing.

### Generation and characterization of singlet oxygen generator-coupled bacteriophages

Gram-positive *L. monocytogenes* specific bacteriophages were coupled with thiorhodamine SOG by mixing 1.2 ml of *Listeria* A500 phage solution (a titer of 10^12^ plaque forming units/ml) with 60 μl of 0.2 M NaHCO_3_ and 30 μl of 0.15 mM NHS-functionalized thiorhodamine. The mixture was incubated at RT in the dark for 1 h. The phage-SOG complex was precipitated by adding 1/5 volume of 20% PEG/2.5M NaCl and incubated for 2 hours at 4 °C. The supernatant was discarded to remove unreacted SOG molecules, and the M13-dye pellet was resuspended in 1 ml of SM buffer and subjected to plaque-forming unit (PFU) determination. For the PFU assay, 990 μl overnight culture of *L. monocytogenes* cells were mixed with 10 μl of serially diluted phage stock solution (in SM buffer). The culture was then mixed and vortexed with 5 ml of melted soft 1/2 BHI agar (50°C) and poured onto the solid agar plates. Solidified agar plates were incubated at 30 °C for a day and phage titer determined based on observed plaque numbers.

### Bacteriophage-guided exploration of virus-targeted host surfaceomes

*Listeria monocytogenes* cells (strain WSLC 1042) were treated with 40 μg/ml gentamicin for 1 hour at RT before incubation with SOG-coupled *Listeria* A500 phages at a multiplicity of infection (MOI) of 10 for 30 min at 4°C in the dark. Per sample, bacteria corresponding to 35 x 10^9^ colony-forming units (CFU) were washed with ice-cold PBS, resuspended in chilled photo-oxidation buffer and illuminated at a working distance of 20 cm with 590-nm Precision LED spotlights for 15 min or left in the dark (three biologically independent samples each). Cells were pelleted by centrifugation and resuspended in chilled labeling buffer for 50 min at 4 °C in the dark. For flow cytometric analysis of cell surface biotinylation, cells were extensively washed with PBS and stained with NeutrAvidin Protein DyLight 488 1:50 for 20 min prior analysis using an Accuri C6 Flow Cytometer (BD Biosciences) with side-scatter (height) set to 10’000 as detection limit and FlowJo (v.10.07). The FSC/SSC pattern of the unlabeled condition was thereby used to gate for live cells. For LUX-MS analysis, cells were extensively washed with PBS, snap-frozen as cell pellets in liquid nitrogen and stored at −80 °C until further processing.

### Genetic engineering and microscopic validation of identified *L. monocytogenes* surface proteins

Candidate genes encoding putative surface proteins A0A0E1RAE1 (LPXTG-motif cell wall anchor domain protein) and A0A0E1R383 (ABC transporter) and their native promoters were cloned into the pPL2 integration vectors^63^ with an HA tag using Gibson method as previously described^86^. The empty vector pPL2, two insert-containing vectors pPL2(*05780*-HA) and pPL2(*06470*-HA), were transformed into electrocompetent *L. monocytogenes* WSLC 1042 cells. The primers 5’-GTCAAAACATACGCTCTTATC-3’ and 5’-ACATAATCAGTCCAAAGTAGATGC-3’ were used to verify the pPL2 integration into the bacterial genome. For surface immunostaining of *L. monocytogenes* WT and HA-tagged strains, corresponding cells were grown to mid-log phase, washed with PBS and treated with 100 μg/ml lysozyme for 30 min to make surface proteins more accessible for immunostaining and then blocked in 1% BSA/PBS. Cells were further washed and stained with an InlA polyclonal antibody (LSBio, Inc) or an HA monoclonal antibody conjugated with Alexa Fluor 555 (ThermoFisher Scientific) at a 1:100 concentration for 30 min prior to visualization by confocal microscopy. For InlA-staining, cells were washed again in PBS and further incubated in Alexa Fluor 488-conjugated goat anti-rabbit secondary antibody. After a final wash step, cell suspensions were applied to a glass slide and imaged by confocal microscopy.

### Synthesis and characterization of singlet oxygen generator-coupled immunogenic peptide gp33

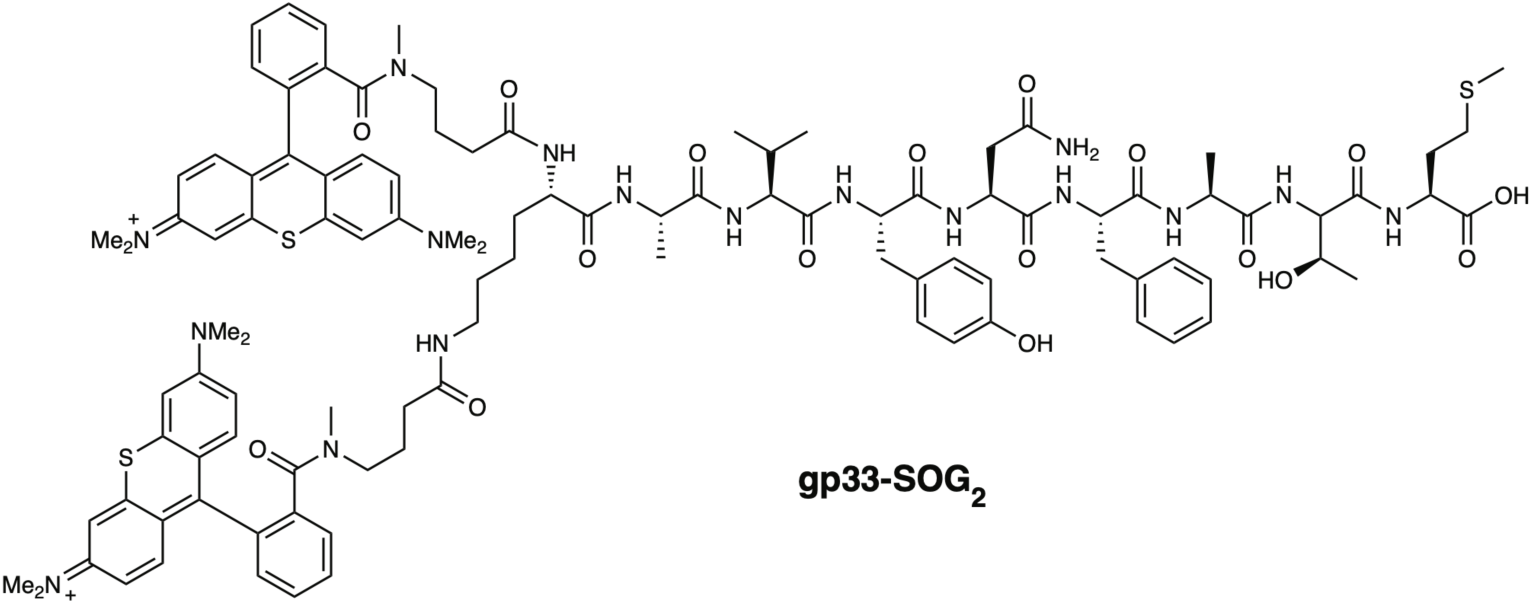

LCMV peptide gp33-41 (gp33; KAVYNFATM) was purchased from NeoMPS. Peptide gp33 (1.0 mg, 1.0 μmol, 1 equiv.) and NHS-thiorhodamine (1.4 mg, 2.0 μmol, 2 equiv.) were incubated at RT for 90 min in PBS at pH 8.3. The reaction mixture was directly purified by preparative reverse-phase high-performance liquid chromatography (RP-HPLC) on a Jasco instrument using a YMC C18 column (5 μm, 20 mm I.D. x 250 mm) at RT at a flow rate of 10 mL/min, with simultaneous monitoring of the eluent at 220, 254 and 301 nm. Milli-Q water with 0.1% TFA (solvent A) and a gradient of 5 to 15% ACN with 0.1% TFA (solvent B) over 5 min and 15 to 75% solvent B over 30 min was used as the eluent. Fractions containing pure product were pooled and lyophilized to obtain gp33-SOG_2_ as a purple solid. The purity and identity were verified by analytical RP-HPLC and HR-MS (**Supplementary Fig. 12**). HR-MS (ESI): calculated for C_106_H_133_N_17_O_17_S_3_^2+^ [M]^2+^: 1005.9608, found: 1005.9589.

### Functional assessment of singlet oxygen generator-coupled gp33

CD8^+^ T cells were isolated from smashed spleens and inguinal lymph nodes of CD45.1 x Nur77-GFP mice using the EasySep(tm) Mouse naïve CD8^+^ T cell Isolation Kit (StemCell, Grenoble, France) following the manufacturer’s instructions. Spleens and lymph nodes were smashed using a 70µm cell strainer. Isotopically labeled mouse dendritic cells were incubated with no compound or 1 μg/ml purified gp33 carrying no (gp33), one (gp33-SOG) or two (gp33-SOG_2_) singlet oxygen generators for 1 h. A 10-fold excess of acutely isolated CD8^+^ T-cells was added by centrifugation and cells were co-incubated for 3 h in the presence of interleukin-2. Cells were carefully washed with ice-cold PBS, chilled photo-oxidation buffer was added and interacting cells were illuminated at a working distance of 20 cm with 590-nm Precision LED spotlights for 15 min. Cells were pelleted by centrifugation and resuspended in chilled labeling buffer for 50 min at 4 °C in the dark. Cells were extensively washed with PBS and assessed for T-cell activation and cell-type specific surface biotinylation using flow cytometry.

### Light-activated *in situ* labeling of functional immunosynapses

Per sample, ten million isotopically labeled mouse dendritic cells were pre-activated with 0.5 µg/ml CpG (Microsynth, Balgach, Switzerland) overnight. PBS-washed dendritic cells were then incubated with 1 μg/ml gp33-SOG_2_ for 1 h in the dark to minimize background light-induced oxidation. A 1.5-fold excess of freshly isolated CD8^+^ T cells was added by centrifugation and cells were co-incubated for 30 min in the presence of interleukin-2. Cells were carefully washed with ice-cold PBS, chilled photo-oxidation buffer was added and interacting cells were illuminated at a working distance of 20 cm with 590-nm Precision LED spotlights for 15 min or left in the dark (two and three biologically independent samples, respectively). Cells were pelleted by centrifugation and resuspended in chilled labeling buffer for 50 min at 4 °C in the dark. Cells were extensively washed with PBS, snap-frozen as cell pellets in liquid nitrogen and stored at −80 °C until further processing.

### Flow cytometry analysis of murine immune cells

Harvested cells were stained for flow cytometry in PBS for 20 min at RT. The following antibodies were purchased from BioLegend (San Diego, United States) and used for flow cytometric analysis: CD8a-BV510 (53-6.7), CD45.1-PacificBlue (A20), CD44-APC (IM7), CD11c-PerCP (N418), CD69-PeCy7 (H1.2F3), CD25-FITC (3C7). Viability of cells was determined by using fixable near-IR dead cell staining (LifeTechnologies, Carlsbad, United States). Biotinylation was assessed by streptavidin-PE (BioLegend) staining for 10 min at room temperature. T cell activation was determined by expression of GFP expression under the control of the Nur77 promoter. Data was acquired on a Canto^™^ flow cytometer (BD Bioscience, Allschwil, Switzerland) and analyzed using the FlowJo software (Treestar, Ashland, OR, USA).

### Automated protein capture and processing

Photo-labeled cell pellets were lysed in 500 μl lysis buffer (100 mM Tris, 1% sodium deoxycholate, 10 mM Tris(2-carboxyethyl)phosphine (TCEP), 15 mM 2-chloroacetamide) by sonication for four intervals of 30 seconds in a VialTweeter (Hielscher Ultrasonics) at a power of 170 W with 80% cycle time and subsequent heating to 100 °C for 5 min. The lysis buffer for bacterial cells additionally contained 75 μg Ply511 *Listeria* endolysins to diminish cell wall integrity. Protein concentration was determined using a Nanodrop 2000 Spectrophotometer (Thermo Fisher Scientific) and equal amounts between sample and control condition (typically 3 mg protein) were subjected to automated capture and processing of photolabeled proteins using a liquid-handling robot. Specifically, in-house packed tips containing 80 μl Streptavidin Plus UltraLink resin (Thermo Fisher Scientific) were hooked to a Versette liquid-handling system (Thermo Fisher Scientific) for automated mixing with cell lysate for 2.5 h and subsequent washing with 5 M NaCl, StimLys buffer (50 mM Tris pH 7.8, 137 mM NaCl, 150 mM glycerol, 0.5 mM EDTA, 0,1% Triton X-100), 100 mM NaHCO_3_ and 50 mM (NH_4_)HCO_3_. Bead-bound proteins were enzymatically digested with 0.5 μg lysyl endopeptidase Lys-C (Wako, cat no. 125-05061) for 2 h at 37°C in 3 M urea / 50 mM ammonium bicarbonate and subsequently diluted to 1.5 M urea / 50 mM ammonium bicarbonate for overnight digestion with 0.8 μg sequencing-grade trypsin at 37°C. Eluting peptides were C18-purified using 5–60 μg UltraMicroSpin Columns according to manufacturer’s instructions and resuspended in 3 % acetonitrile (ACN), 0.1% formic acid (FA) containing iRT peptides (Biognosys AG, Schlieren, Switzerland) for mass spectrometric analysis.

### Liquid chromatography-tandem mass spectrometry (LC-MS/MS)

For antibody-, small molecule-, biomolecule- and Thanatin-guided LUX-MS samples acquired in data-dependent acquisition (DDA) mode, peptides were separated by reversed-phase chromatography on an HPLC column (75-μm inner diameter, New Objective) that was packed in-house with a 15-cm stationary phase (ReproSil-Pur C18-AQ, 1.9 μm) and connected to a nano-flow HPLC with an autosampler (EASY-nLC 1000, Thermo Scientific). The HPLC was coupled to a Q-Exactive Plus mass spectrometer (Thermo Scientific) equipped with a nano electrospray ion source (Thermo Scientific). Peptides were loaded onto the column with 100% buffer A (99.9% H_2_O, 0.1% FA) and eluted at a constant flow rate of 300 nl/min with a 70-min linear gradient from 6–28% buffer B (99.9% ACN, 0.1% FA) followed by a 4-min transition from 28 to 50% buffer B. After the gradient, the column was washed for 10 min with 98% buffer B, 4 min with 10% buffer B, and 8 min with 98% buffer B. Electrospray voltage was set to 2.2 kV and capillary temperature to 250 °C. A high-resolution survey mass spectrum (from 300 to 1,700 m/z) acquired in the Orbitrap with resolution of 70,000 at m/z 200 (automatic gain control target value 3 × 10^6^) was followed by MS/MS spectra in the Orbitrap at resolution of 35,000 (automatic gain control target value 1 × 10^6^) of the 12 most-abundant peptide ions with a minimum intensity of 2.5 × 10^4^ that were selected for fragmentation by higher-energy collision-induced dissociation with a collision energy of 28% and an isolation window of 1.4 Da. Fragmented precursors were dynamically excluded for 30 s.

For bacteriophage-guided LUX-MS samples acquired in data-dependent acquisition (DDA) mode, peptides were separated by reversed-phase chromatography on an HPLC column (75-μm inner diameter, New Objective) that was packed in-house with a 15-cm stationary phase (ReproSil-Pur C18-AQ, 1.9 μm) and connected to a nano-flow HPLC with an autosampler (EASY-nLC 1000, Thermo Scientific). The HPLC was coupled to a Orbitrap Fusion Tribrid mass spectrometer (Thermo Scientific) equipped with a nano electrospray ion source (Thermo Scientific). Peptides were loaded onto the column with 100% buffer A (99.9% H_2_O, 0.1% FA) and eluted at a constant flow rate of 300 nl/min with a 70-min linear gradient from 6–28% buffer B (99.9% ACN, 0.1% FA) followed by a 4-min transition from 28 to 50% buffer B. After the gradient, the column was washed for 10 min with 98%, 4 min with 10% and again 8 min with 98% buffer B. Electrospray voltage was set to 1.8 kV and capillary temperature to 275. A high-resolution survey mass spectrum (from 395 to 1,500 m/z) acquired in the Orbitrap with resolution of 120,000 at m/z 200 (automatic gain control target value 2 × 10^5^ and maximum injection time 50 ms) was followed by MS/MS spectra of most-abundant peptide ions with a minimum intensity of 5 × 10^3^ that were selected for subsequent higher-energy collision-induced dissociation fragmentation with a collision energy of 35% and an isolation window of 1.6 Da. Fragments were detected by MS/MS acquisition in the Ion Trap with scan rate set to “Rapid” (automatic gain control target value 1 × 10^4^ and maximum injection time 250 ms). Fragmented precursors were dynamically excluded for 30 s.

For antibody-guided LUX-MS samples acquired in data-independent acquisition (DIA) mode, peptides were separated by reversed-phase chromatography on an HPLC column (75-μm inner diameter, New Objective) that was packed in-house with a 15-cm stationary phase (ReproSil-Pur C18-AQ, 1.9 μm) and connected to a nano-flow HPLC with an autosampler (EASY-nLC 1200, Thermo Scientific). The HPLC was coupled to a Orbitrap Fusion Tribrid mass spectrometer (Thermo Scientific) equipped with a nano electrospray ion source (Thermo Scientific). Peptides were loaded onto the column with 100% buffer A (99.9% H_2_O, 0.1% FA) and eluted at a constant flow rate of 250 nl/min with a 70-min linear gradient from 6–28% buffer B (99.9% ACN, 0.1% FA) followed by a 4-min transition from 28 to 50% buffer B. After the gradient, the column was washed for 10 min with 98% buffer B, 4 min with 10% buffer B, and 8 min with 98% buffer B. Electrospray voltage was set to 4.0 kV and capillary temperature to 320 °C. A survey scan acquired in the orbitrap with 120,000 resolution, 50 ms max injection time, and an AGC target of 3×10^6^ was followed by 24 precursor windows with fragmentation at a collision energy of 28% and MS/MS spectra acquisition in the orbitrap with 30,000 resolution. The injection time was set to auto, AGC target to 1×10^6^ and mass range to 300-1700 m/z. For the spectral library, the samples were additionally measured on the Q-Exactive Plus platform in data-dependent acquisition (DDA) mode as described above

For gp33-guided LUX-MS samples acquired in data-independent acquisition (DIA) mode, peptides were separated by reversed-phase chromatography on an HPLC column (75-μm inner diameter, New Objective) that was packed in-house with a 30-cm stationary phase (ReproSil-Pur C18-AQ, 1.9 μm) and connected to a nano-flow HPLC with an autosampler (EASY-nLC 1200, Thermo Scientific). The HPLC was coupled to a Orbitrap Fusion Lumos Tribrid mass spectrometer (Thermo Scientific) equipped with a nano electrospray ion source (Thermo Scientific). Peptides were loaded onto the column with 100% buffer A (99.9% H_2_O, 0.1% FA) and eluted at a constant flow rate of 250 nl/min with a 240-min non-linear gradient from 4–58% buffer B (80% ACN, 0.1% FA). After the gradient, the column was washed for 10 min with 98% buffer B, 4 min with 1% buffer B. Electrospray voltage was set to 3.6 kV and capillary temperature to 320 °C. A survey scan acquired in the orbitrap with 120,000 resolution, 50 ms max injection time, and an AGC target of 4×10^5^ was followed by 80 precursor windows with fragmentation at a collision energy of 27% and MS/MS spectra acquisition in the orbitrap with 30,000 resolution. The injection time was set to auto, AGC target to 1×10^6^ and mass range to 350-2000 m/z. For the spectral library, replicates of the illuminated and non-illuminated condition were pooled and analyzed on the same instrumental setup in data-dependent acquisition (DDA) mode using a universal method with equal gradient and a collision energy of 27% and 30%.

## Data analysis

Raw files acquired in DDA mode were searched against corresponding SwissProt-reviewed protein databases containing common contaminants using Comet (v.2015.01) within the Trans Proteomic Pipeline v.4.7 (SPC/ISB Seattle). Peptides were required to be fully tryptic with a maximum of two missed cleavage sites, carbamidomethylation as fixed modification, and methionine oxidation as a dynamic modification. The precursor and fragment mass tolerance was set to 20 ppm and 1 Da (MS2 ion trap) or 0.02 Da (MS2 orbitrap), respectively. Proteins identified by at least two proteotypic peptides were quantified by integration of chromatographic traces of peptides using Progenesis QI v.4.0 (Nonlinear Dynamics). Contaminant hits were removed, and proteins filtered to obtain a false discovery rate of < 1%. Raw protein abundances were exported based on non-conflicting peptides.

For DIA analysis, raw files for spectral libraries acquired in DDA mode were searched against corresponding SwissProt-reviewed protein databases containing common contaminants using the Sequest HT search engine within Thermo Proteome Discoverer version 2.4 (Thermo Scientific). Peptides were required to be fully tryptic with up to two missed cleavages. Carbamidomethylation was set as a fixed modification for cysteine. Oxidation of methionine and N-terminal acetylation were set as variable modifications. For SILAC data, isotopically heavy labeled arginine (+10 Da) and lysine (+8 Da) were set as variable modifications. Monoisotopic peptide tolerance was set to 10 ppm, and fragment mass tolerance was set to 1 Da (MS2 ion trap) or 0.02 Da (MS2 orbitrap). The identified proteins were assessed using Percolator and filtered using the high peptide confidence setting in Protein Discoverer. Analysis results were then imported to Spectronaut v.13 (Biognosys AG, Schlieren, Switzerland) for the generation of spectral libraries filtering for high confidence identification on peptide and protein level. Raw files acquired in data-independent acquisition mode were analyzed using Spectronaut v.13 with default settings. The proteotypicity filter “only protein group specific” was applied, and extracted feature quantities were exported from Spectronaut using the “Quantification Data Filtering” option. Within the R computing environment (v.3.4.0), protein abundance changes (expressed in log2) were calculated using a linear mixed effect model and tested for statistical significance using a two-sided t-test in the R package MSstats (v.3.8.6) ^85^. For human proteins, cell surface localization was inferred by matching protein identifications to the in silico human surfaceome ^11^. For all other organisms, cell surface localization was inferred based on UniProt GO-term annotations being either “cell surface”, “cell membrane” or “secreted”. Significantly enriched proteins (abundance fold change > 1.5 and p-value < 0.05) were considered as acute proximity candidates of the ligand- or antibody-SOG construct. The CD20 surfaceome interaction network was generated using Cytoscape (v.3.7.1)^87^. Protein topology information was retrieved and visualized using the PROTTER interactive webtool^88^. Protein structures were visualized using PyMOL Molecular Graphics System (v.2.4.0, Schrödinger, LLC.). Schematic figures were created using the BioRender web tool (https://biorender.com/). Interactive volcano plots were created using an in-house developed R script using the plotly package^89^.

## Data availability

Source data for **Fig. 2d-f** and **Supplementary Fig. 5** are provided in the paper in **Supplementary Table 1**. Source data for **Fig. 3c** and e are provided in the paper in **Supplementary Table 2**. Source data for **Fig. 4b** are provided in the paper in **Supplementary Table 3**. Source data for **Fig. 5c** are provided in the paper in **Supplementary Table 4**. Source data for **Fig. 6c-d** are provided in the paper in **Supplementary Table 5**. The original mass spectra have been deposited to the ProteomeXchange Consortium (http://proteomecentral.proteomexchange.org) via the PRIDE partner repository^90^ with the dataset identifier PXD020481. Any additional data that support the findings of this study are available from the corresponding author upon reasonable request.

## Acknowledgements

We acknowledge Julia Boshard, Sebastian Steiner and Dr. Patrick Pedrioli for constructive discussions and specific comments on the manuscript. We specifically acknowledge Dr. Damaris Bausch-Fluck and Dr. Andreas Frei for their strategic input and critical discussions in the conceptional stage of the project. We are grateful to the members of the B.W. research group for suggestions and support at all stages of the project. We thank J. R. Wyatt for scientific text editing. We thank Stephan Schneider for the technical assistance on phage enrichment and coupling with SOGs. We further acknowledge Dr. Sebastian Müller, Dr. Patrick Pedrioli and Dr. Silvana Albert for assistance in analysing the SILAC-DIA LUX-MS dataset and David Vogel for contributing to antibody-guided LUX-MS experiments. The following agencies contributed and are acknowledged for their generous support: ETH (grant ETH-30 17-1 and grant ETH-25 15-2) and Swiss National Science Foundation (grant 31003A_160259) for B.W. and NIH grants R01-GM-094231 and U24-CA210967 for A.I.N.

## Author contributions

M.M. performed all experiments except those noted below. F.G. and N.B. performed isolation and FACS analysis of murine immune cells. Y.S. produced SOG-coupled bacteriophages and performed immunofluorescence experiments. S.U.V. and M.Mo. synthesized SOG-coupled Thanatin. J.R.P. provided CG1 and scientific expertise. Y.Se. performed isolation of PBMC. M.v.O. contributed to method optimization. R.H. and R.S. synthesized SOG-coupled gp33. A.I.N contributed new analytical tools. M.M. and B.W. optimised and performed LC-MS/MS acquisition. M.M., F.G. and N.B. analyzed data. M.M., F.G., N.B., Y.S., J.R.P., E.C., J.B., B.S., J.A.R., M.J.L., A.O. and B.W. designed research. M.M. and B.W. conceived the project and wrote the paper. All authors reviewed and commented on the manuscript.

---

**Supplementary Figure 1.**
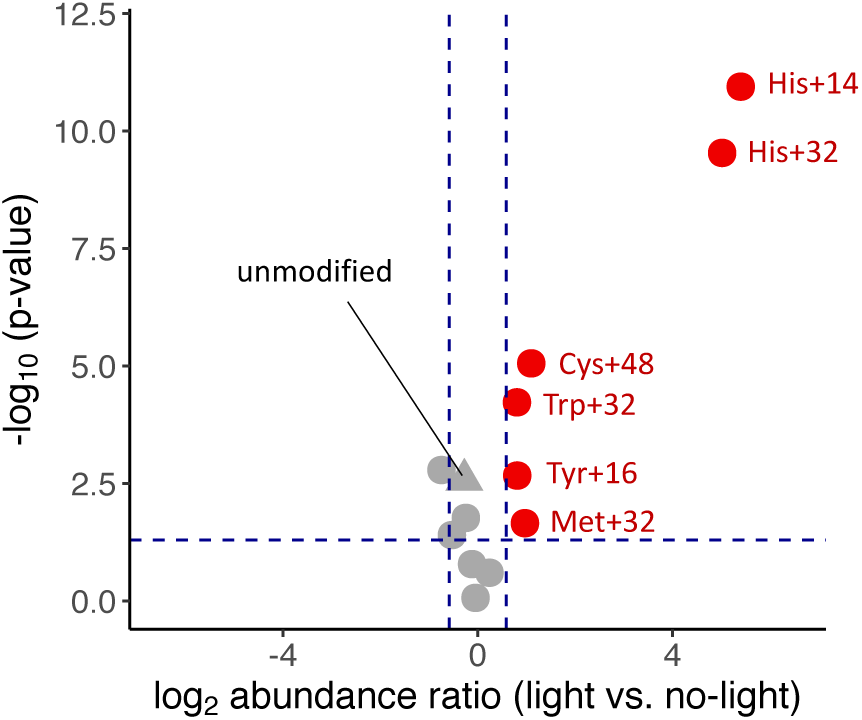
Identification and quantification of light-induced protein modifications. Volcano plot showing relative abundance of transferrin proteins with indicated amino acid modifications as quantified by mass spectrometry before and after illumination for 15 min. Dots and triangles represent modified and unmodified transferrin proteins, respectively. Grey and red indicates non-regulated and significantly regulated proteins.

**Supplementary Figure 2.**
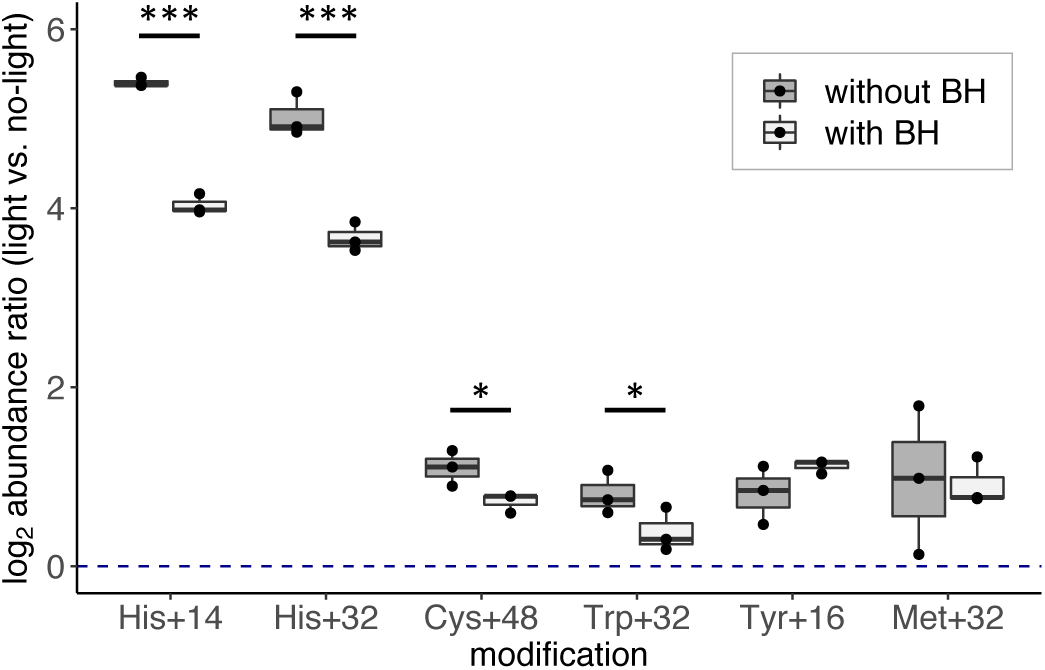
Hydrazide-linker reactivity of light-induced protein modifications. Barplot showing quantitative generation of modified transferrin proteins after illumination for 15 min in the absence or presence of biocytin-hydrazide (BH) linker. The center line of box-plots represents the median, box limits the upper and lower quartiles, whiskers the 1.5x interquartile range. Statistical analysis was done with two-tailed unpaired Student’s t test. One star indicates P < 0.1, two stars P < 0.01 and three stars P < 0.005.

**Supplementary Figure 3.**
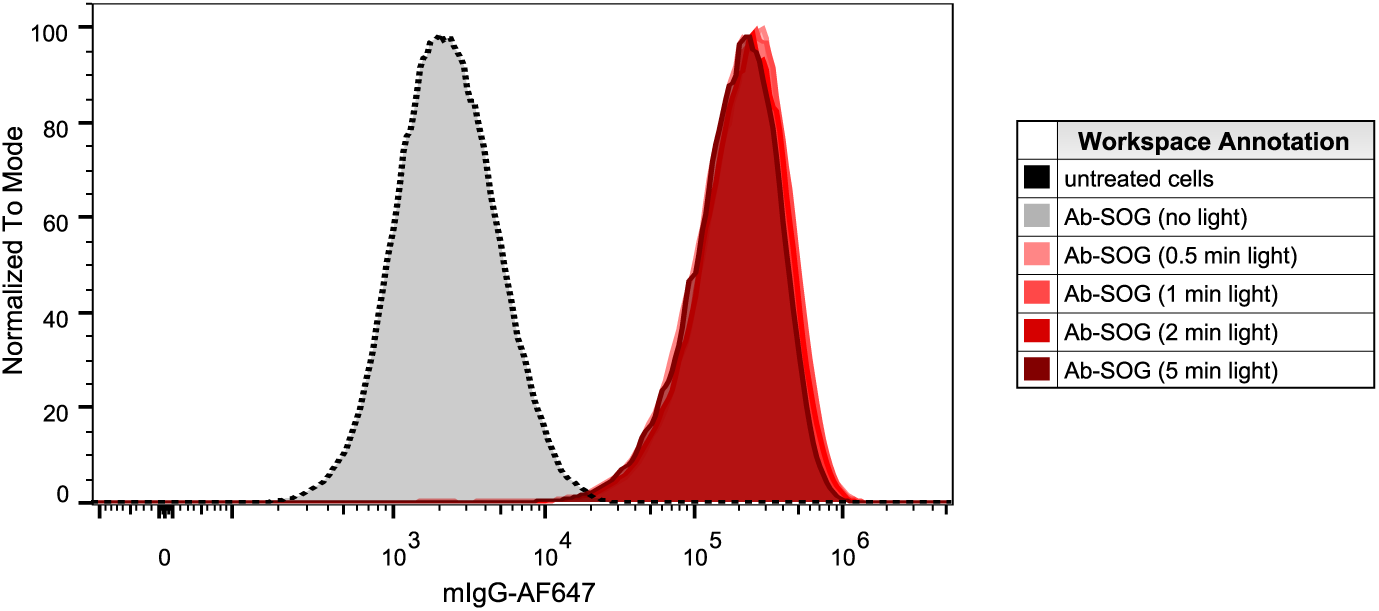
Antibody-SOG construct binding to living cells. Histogram plot showing binding of singlet oxygen generator coupled anti-CD20 antibodies to B-lymphoma SUDHL6 cells that were subsequently illuminated for 0 to 5 min.

**Supplementary Figure 4.**
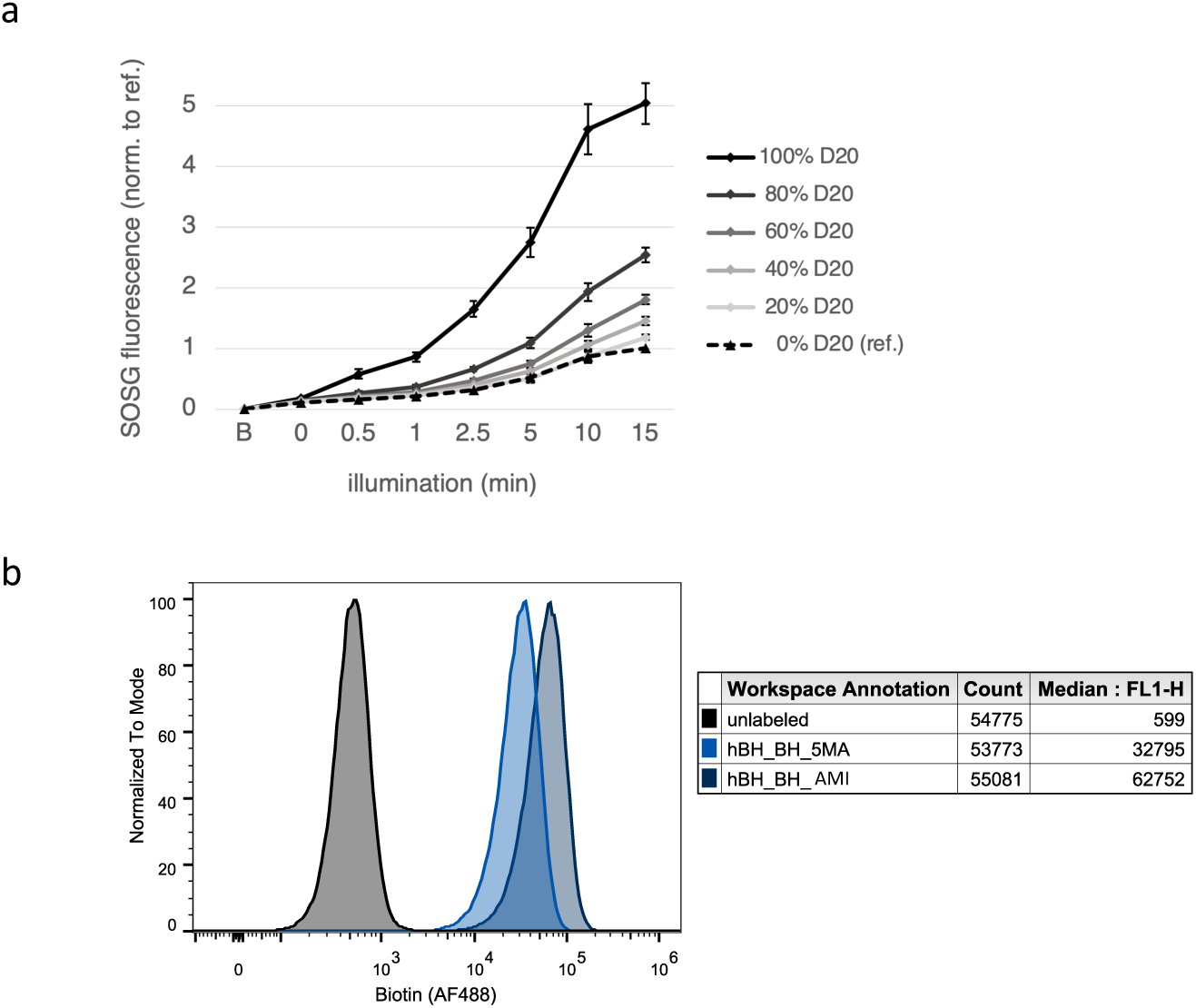
Buffer and catalyst-based fIne-tuning of light-induced cell surface biotinylation. **a**, Profile plots showing light-dependent fluorescence of oxidized singlet oxygen sensor green (SOSG) at increasing D_2_O/H_2_O ratios as read-out for photo-oxidation efficiency. **b**, Histogram plot showing cell surface biotinylation of B-lymphoma SUDHL6 cells that were LUX-labeled in the presence of either 2-Amino-5-methoxybenzoic acid (5-MA) or 2-(Aminomethyl)imidazole dihydrochloride (AMI) to catalyze hydrazone formation between the biocytin-hydrazide linker and photo-oxidized protein residues. The total cell count and median fluorescence of AF488 (FL1-H) is shown.

**Supplementary Figure 5.**
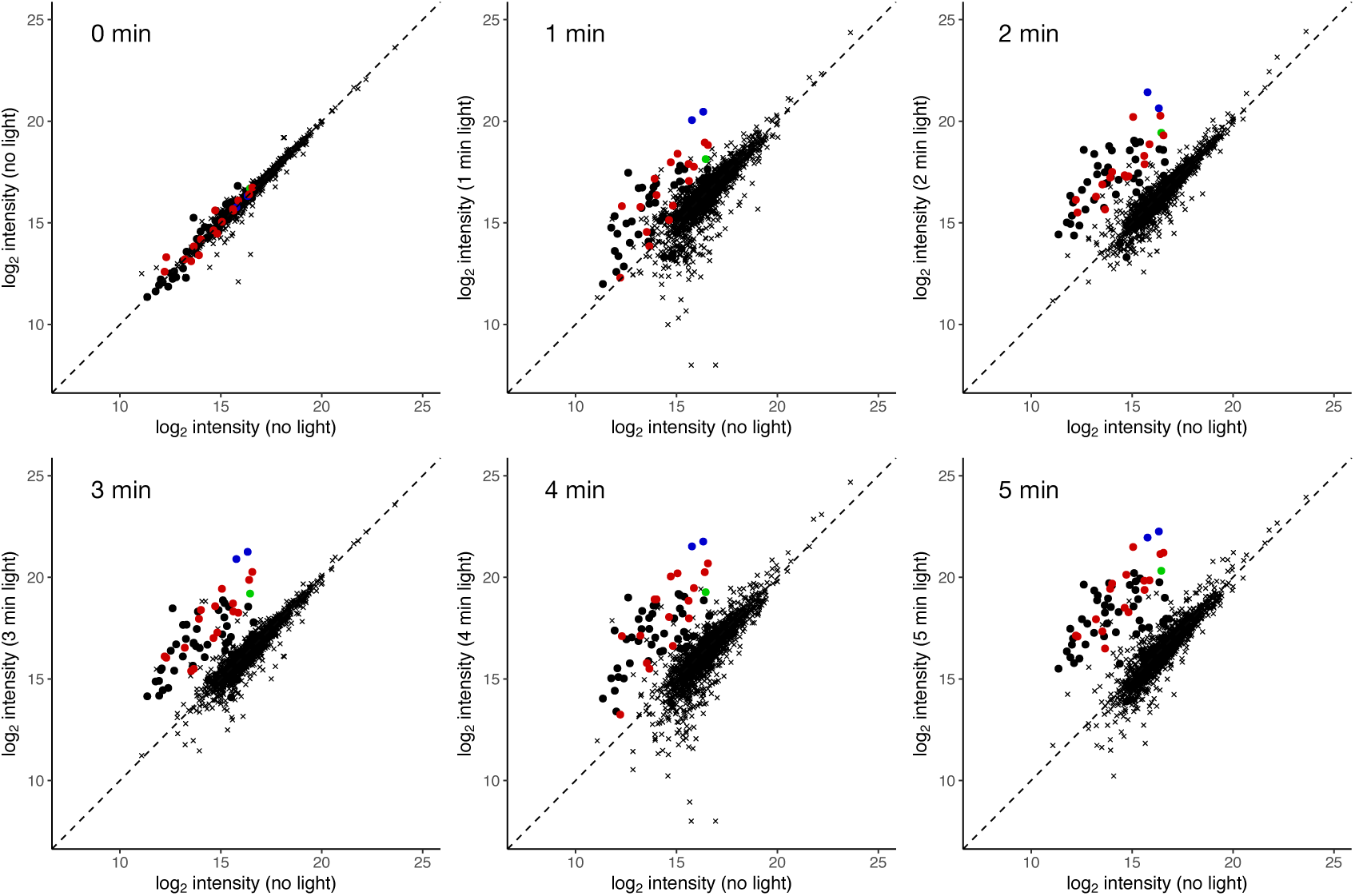
Illumination-dependent fIne-tuning of cell surface biotinylation. **a**, Scatter plots showing relative abundance of LUX-MS quantified proteins from anti-CD20 Ab-SOG treated B-lymphoma SUDHL6 cells between a non-illuminated control (x-axis) or samples illuminated for 0 - 5 min (y-axis). Dots and crosses represent cell surface and otherwise annotated proteins, respectively. Blue, green and red indicate chains of the anti-CD20 antibody, the primary binding target CD20 and known CD20 associated surface proteins.

**Supplementary Figure 6.**
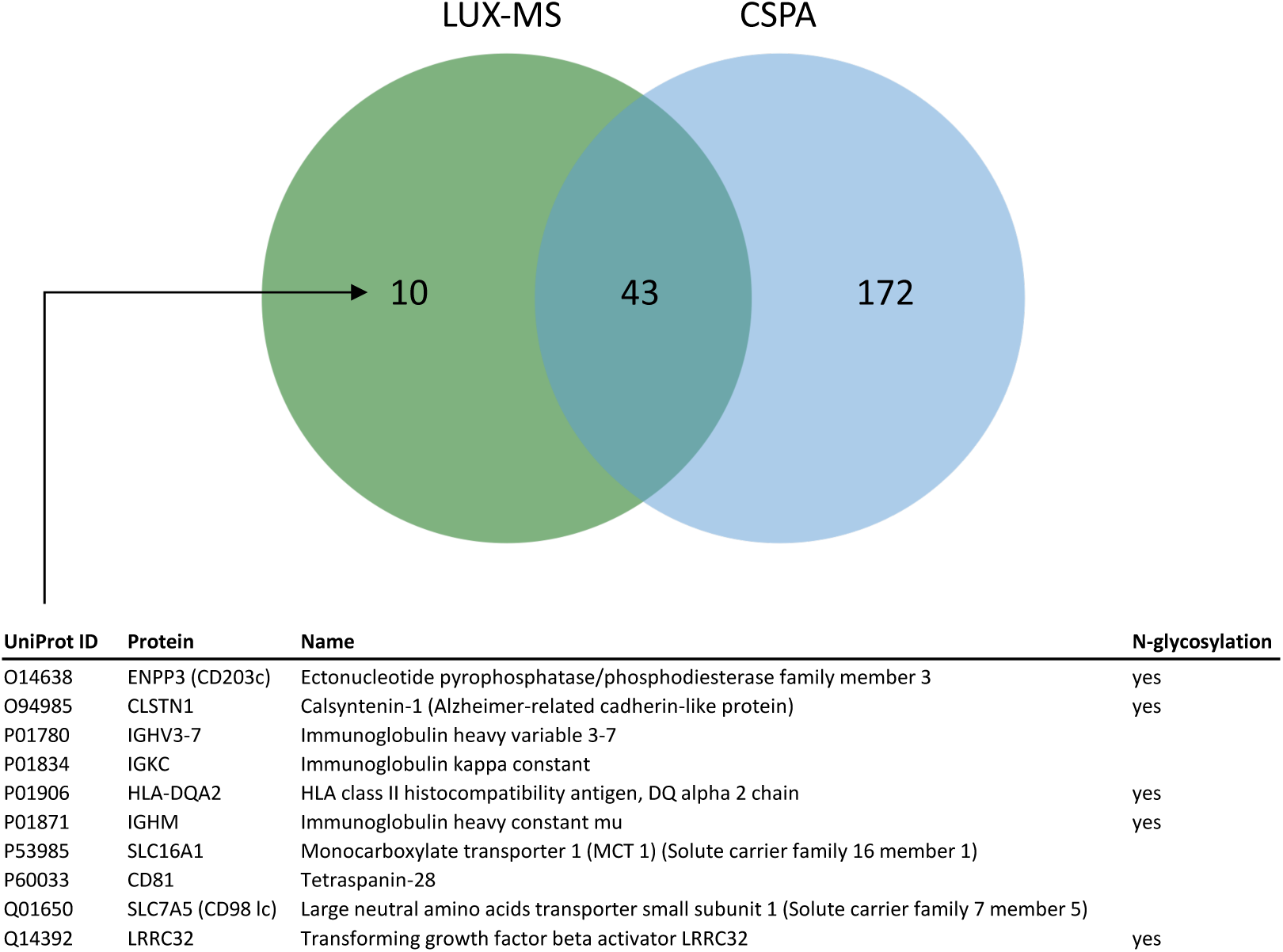
Surfaceome detection by LUX-MS is not restricted to protein N-glycosylation. Venn diagram showing overlap of cell surface proteins identified by anti-CD20 antibody-guided LUX-MS and Cell Surface Capture as reported by the Cell Surface Protein Atlas (CSPA, https://wlab.ethz.ch/cspa/) on B-lymphoma SUDHL6 cells. Cell surface proteins specifically identified by LUX-MS are shown with additional information such as N-glycosylation state.

**Supplementary Figure 7.**
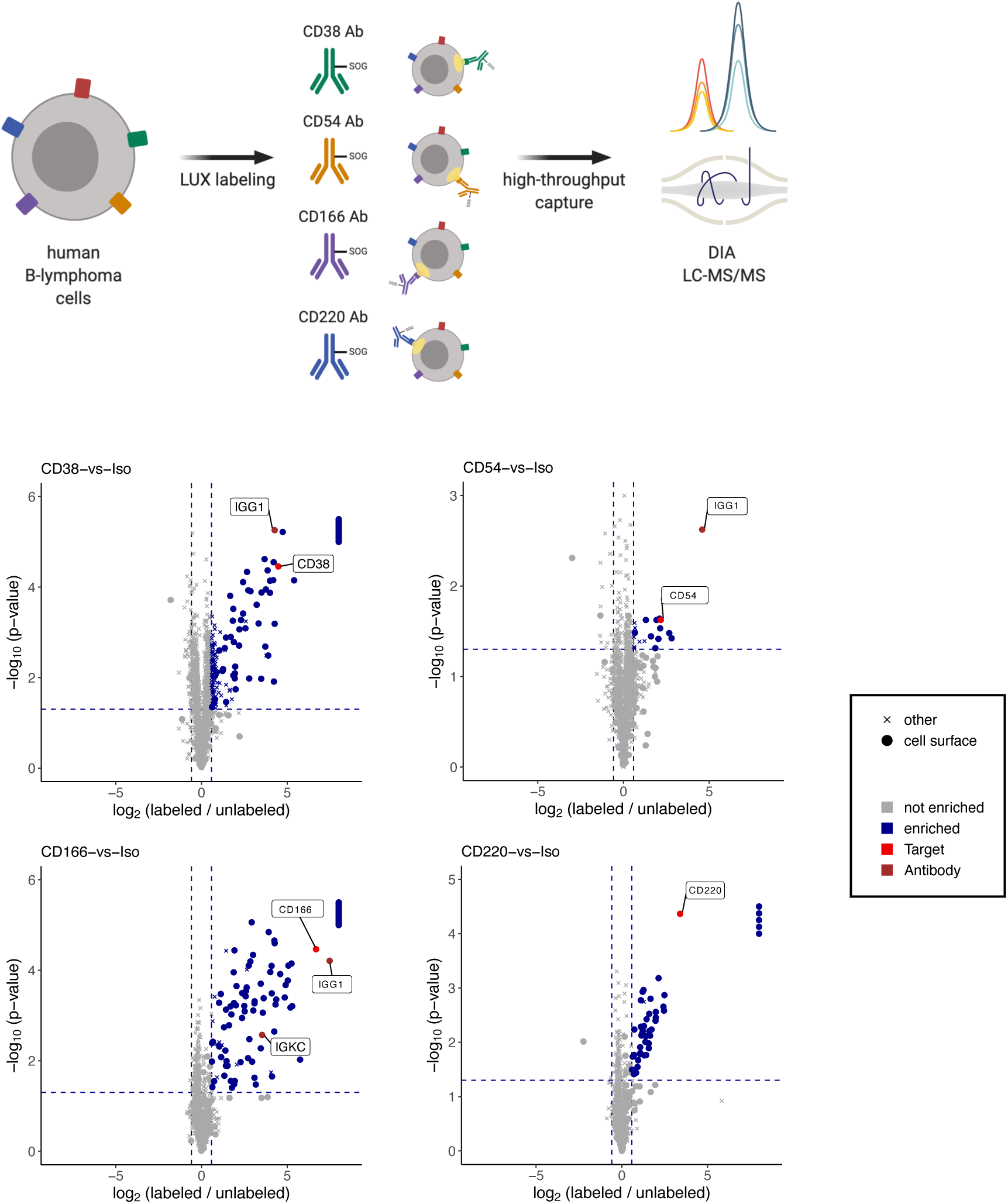
Elucidation of distinct surfaceome receptor neighborhoods on living cells. **a**, Schematic outline of mapping distinct surfaceome receptor neighborhoods on living cells using a selection of singlet oxygen generator (SOG)-coupled antibodies and data-independent acquisition (DIA) mass spectrometry. **b**, Volcano plots showing relative abundance of LUX-MS quantified proteins from B-lymphoma SUDHL6 cells treated with either SOG-coupled isotype control (Iso), anti-CD38, anti-CD54, anti-CD166 or anti-CD220 antibodies and illuminated for 15 min. Dots and crosses represent cell surface and otherwise annotated proteins, respectively. Brown, red and blue indicate chains of the used antibody, the primary binding target and cell surface proximal candidates.

**Supplementary Figure 8.**
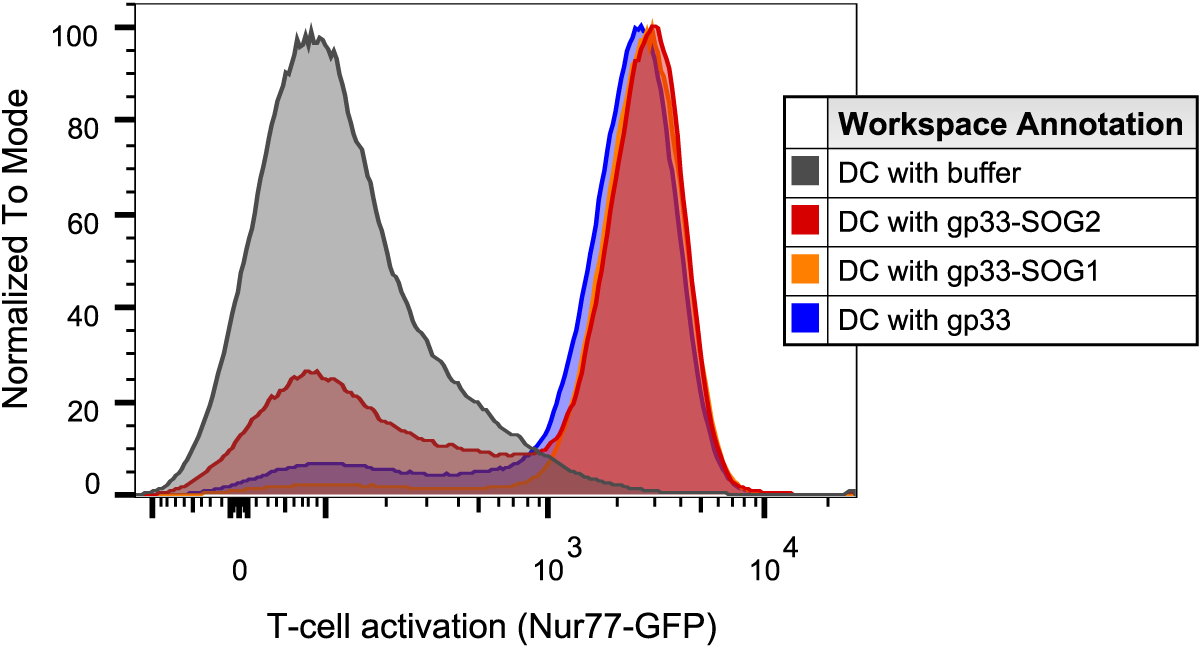
Singlet oxygen generators (SOG) coupling does not affect immunogenicity of gp33. Histogram plot showing activation-indicating expression of Nur77-GFP of gp33-reactive CD8^+^ T-cells after co-culture for 3 h with dendritic cells loaded with either buffer, underivatized gp33, gp33 with one SOG (gp33-SOG1) or gp33 with two SOG (gp33-SOG2).

**Supplementary Figure 9.**
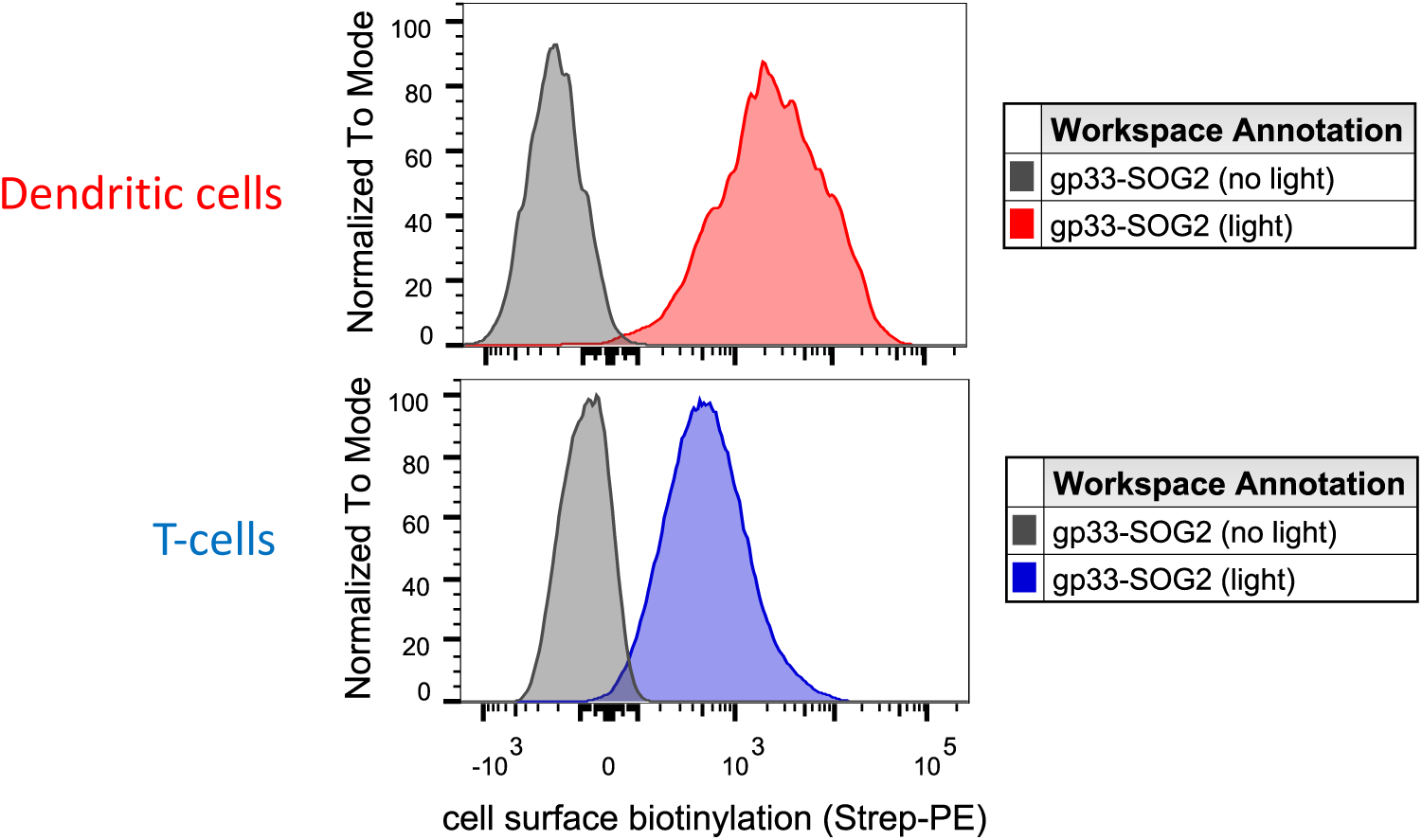
Light-activated proximity labeling of dynamic immune cell interactions. Histogram plots showing cell surface biotinylation of gp33-SOG2 presenting dendritic cells and CD8^+^ T-cells that were co-cultured for 3 h and LUX-labeled with or without illumination for 15 min.

**Supplementary Figure 10.**
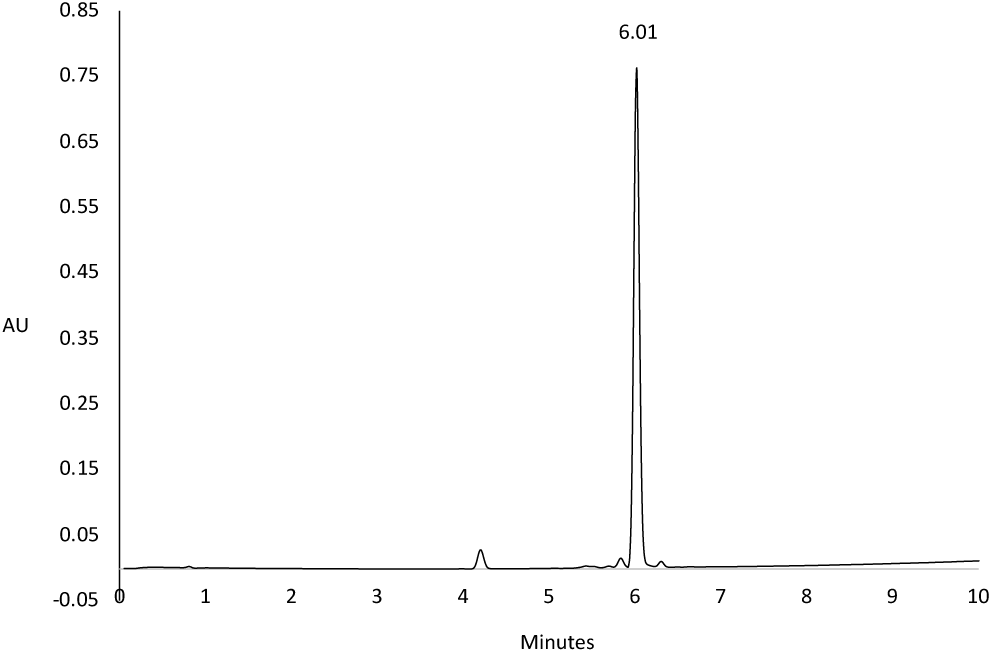
Characterization of CG1-SOG. Analytical RP-HPLC of purified CG1-SOG. Analytical RP-HPLC was performed on a Acquity UPLC C18 column (1.7 μm, 2.1 mm I.D. x 150 mm) at 40 °C at a flow rate of 0.3 mL/min and a gradient of 5 - 95 % solvent B (ACN/H_2_O + 0.1 % TFA) for 10 min.

**Supplementary Figure 11.**
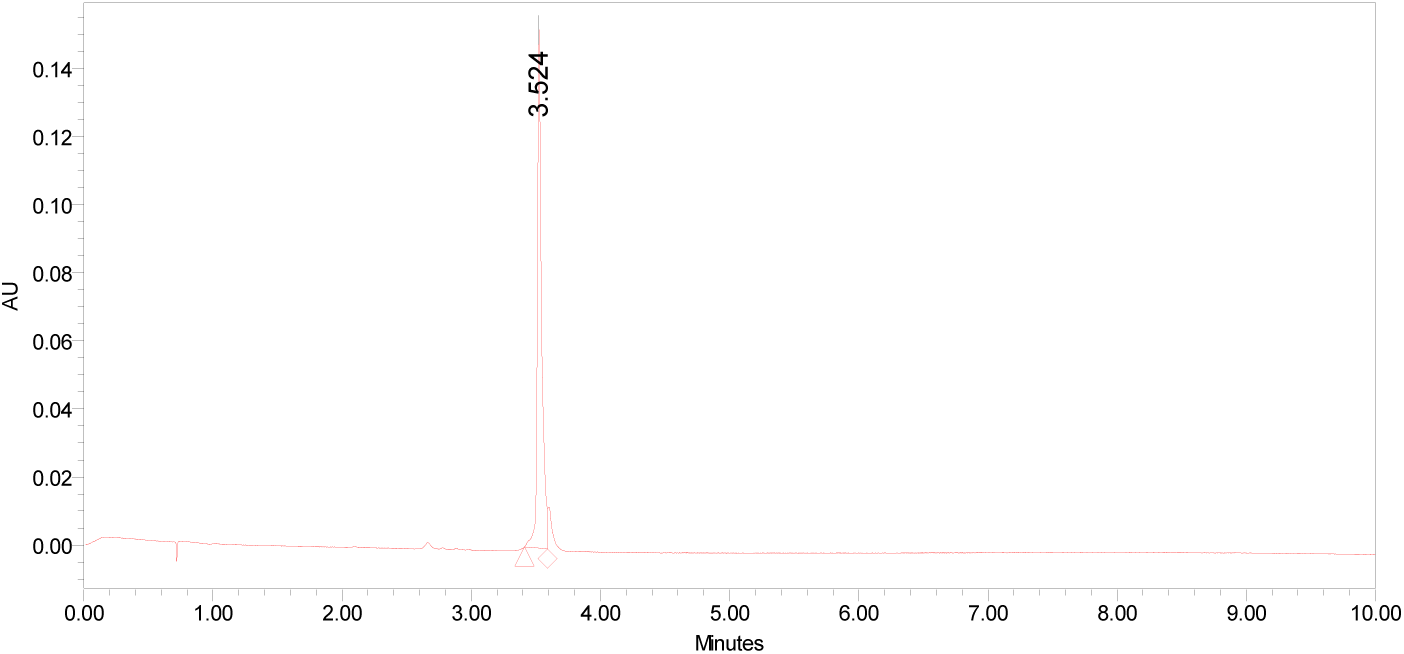
Characterization of Thanatin-SOG. Analytical RP-HPLC of purified Thanatin-SOG. Analytical RP-HPLC was performed on a Acquity UPLC C18 column at 40 °C and a gradient of 5 - 95 % solvent B (ACN/H_2_O + 0.1 % TFA) for 10 min.

**Supplementary Figure 12.**
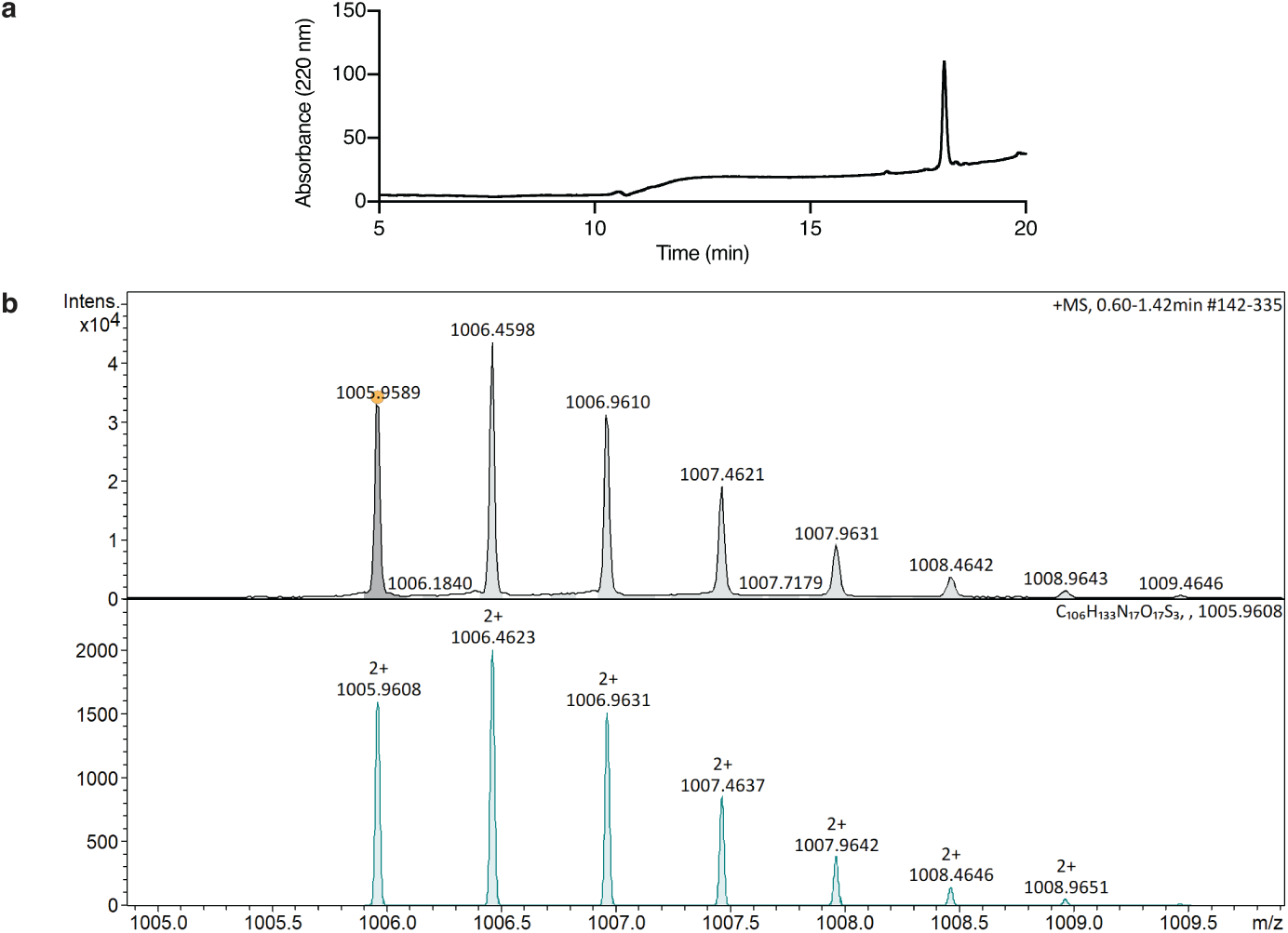
Characterization of gp33-SOG_2_. **a**, Analytical RP-HPLC of purified gp33-SOG_2_. Analytical RP-HPLC was performed on a Shiseido C18 column (5 μm, 4.6 mm I.D. x 250 mm) at RT at a flow rate of 1 mL/min and a gradient of 10% solvent B for 3 min, 10 to 95% solvent B over 14 min, and 95% solvent B for 3 min. **b**, Observed (upper panel) and calculated (lower panel) HR-MS spectra. Spectra were obtained by the mass spectrometry service of the ETH Laboratory of Organic Chemistry on a Bruker Daltonics maXis ESI-QTOF spectrometer.

**Supplementary Table 1** Antibody-guided LUX-MS proteomic data

**Supplementary Table 2** Small molecule drug and biomolecule-guided LUX-MS proteomic data

**Supplementary Table 3** Thanatin-guided LUX-MS proteomic data

**Supplementary Table 4** Bacteriophage-guided LUX-MS proteomic data

**Supplementary Table 5** Immunosynapse LUX-MS proteomic data

**Supplementary Data 1** Interactive volcano plot of anti-CD20 antibody-guided LUX-MS

**Supplementary Data 2** Interactive volcano plot of small molecule drug guided LUX-MS

**Supplementary Data 3** Interactive volcano plot of biomolecule-guided LUX-MS

**Supplementary Data 4** Interactive volcano plot of Thanatin-guided LUX-MS

**Supplementary Data 5** Interactive volcano plot of Bacteriophage-guided LUX-MS

**Supplementary Data 6** Interactive volcano plot of Immunosynapse LUX-MS (dendritic cell proteins)

**Supplementary Data 7** Interactive volcano plot of Immunosynapse LUX-MS (T cell proteins)

